# Conformational asymmetry of replicated human chromosomes

**DOI:** 10.1101/2025.07.09.663929

**Authors:** Flavia Corsi, Sofia Kolesnikova, Thomas L. Steinacker, Zsuzsanna Takács, Paul Batty, Michael Mitter, Daniel W. Gerlich, Anton Goloborodko

**Author notes:** Correspondence, DG, AG. These authors contributed equally to this work (Sofia Kolesnikova, Thomas L. Steinacker). Institut Curie, PSL Research University, Sorbonne Université, CNRS UMR3664, Laboratoire Dynamique du Noyau, 75005 Paris, France. St Anna Children’s Cancer Research Institute (CCRI), 1090 Vienna, Austria. CeMM - Research Center for Molecular Medicine of the Austrian Academy of Sciences, 1090 Vienna, Austria (Paul Batty).

## Abstract

DNA replication creates two sister chromatids that must acquire specific three-dimensional conformations to support genome function and stability. This organization is largely mediated by cohesin complexes, which extrude intra-chromosomal loops and link two chromatids, thus forming “chromatid cohesion”. Although sister chromatids are genetically identical, the replication process is intrinsically asymmetric: each chromatid inherits a different parental DNA strand, while the new strands are synthesized using distinct “leading” and “lagging” mechanisms of the replication fork. Whether and how this molecular asymmetry impacts higher-order chromatin organization remains unknown. Using sister-chromatid-sensitive Hi-C, strand-specific FISH, and polymer modeling, we reveal a consistent, genome-wide shift in sister chromatid alignment, biased along the 5′-3′ direction of the inherited strands. This shift persists without loop extrusion but is lost upon disruption of cohesion, implicating cohesive cohesins in maintaining the displacement. Polymer simulations indicate that a modest (∼100 kb) misalignment of “cohesive” cohesins is responsible for the observed asymmetry. We propose two mechanistic models that explain how this displacement arises from replication fork asymmetry: either through the dislocation of cohesin during replication or through the asymmetric anchoring and subsequent random sliding of cohesin pairs. These findings reveal a previously unrecognized chromosome-scale asymmetry in sister chromatid organization, which has implications for homology search during DNA repair.

## Introduction

After DNA replication, the two newly formed sister chromatids must acquire specific three-dimensional conformations essential for their function and stability. Central to this organization are ring-shaped cohesin SMC complexes that perform two structurally and functionally distinct roles. “Extrusive” cohesins dynamically form DNA loops through their intrinsic motor activity (*1–5*), creating topologically associating domains (TADs) (*6–8*) and facilitating long-range enhancer-promoter communication (*9*). “Cohesive” cohesins link sister chromatids from the S phase until mitosis by entrapping both DNA molecules (*10–13*). The resulting connections and alignment between sisters are essential for their correct segregation in mitosis and homology-directed repair (*14–16*).

While replication produces genetically identical chromatids, its mechanism is inherently asymmetric in two distinct ways (*17*). At the chromosomal level, two chromatids created by semi-conservative replication inherit different parental DNA strands with opposite directionality (Fig. 1A) (*18*). At the molecular level, replication is performed by pairs of opposing asymmetric replication forks (Fig. 1B). These forks comprise a CMG helicase that tracks along only one of the two parental strands (Fig. 1B) (*19, 20*), two distinct sets of enzymes that synthesize “leading” and “lagging” strands, and accessory proteins (*21*). As these replication forks progress, they interact with cohesin complexes, establishing the cohesive linkages that will hold sister chromatids together (*12, 13, 22–29*). Whether and how the asymmetry of the replication process affects the establishment of loops and cohesive linkages remains unknown.

**Figure 1.**
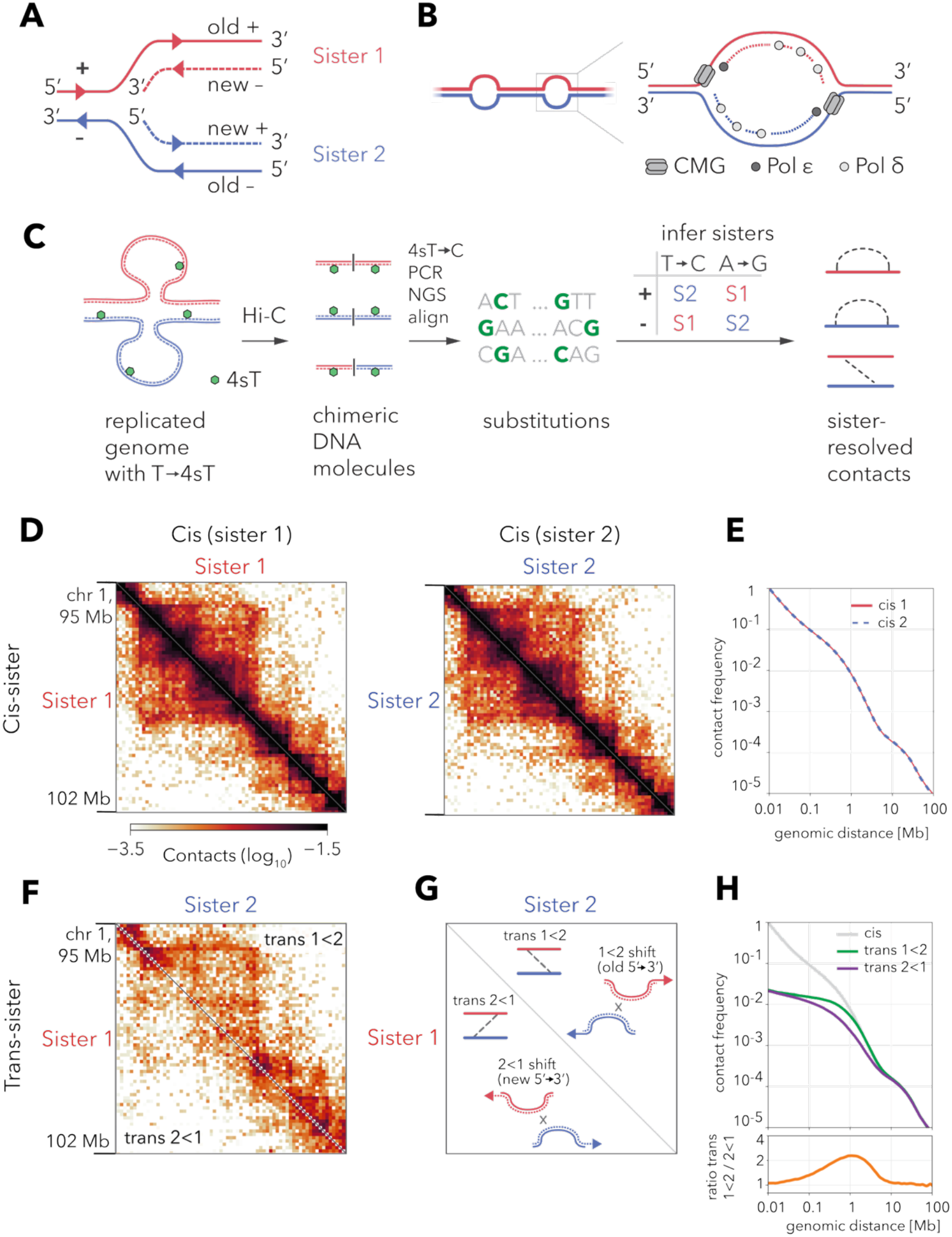
Sister-chromatid-sensitive Hi-C uncovers an asymmetric shift of sister chromatids. (A) Cartoon illustrating semi-conservative DNA replication: Sister 1 (red) inherits the parental (old) + strand, Sister 2 (blue) the – strand. (B) Left: The genome is replicated by bidirectional asymmetric replication bubbles. Right: Within each bubble, the CMG helicase moves 3′→5′ along the leading strand template. Pol ε synthesizes the leading strand continuously while Pol δ synthesizes the lagging strand discontinuously, creating molecular asymmetry at the fork. (C) Workflow of sister-chromatid-sensitive Hi-C (scsHi-C). Newly synthesized DNA is pulse-labeled with 4-thiothymidine (4sT, green), which is converted to T→C substitutions during library preparation, and sequenced. The strand on which the substitution occurs assigns each read to Sister 1 or Sister 2, yielding sister-resolved contact pairs. (D) Cis-sister scsHi-C contact maps at 100 kb resolution for a 7 Mb window on chromosome 1, Sister 1 (left), and Sister 2 (right). (E) Plots of cis-sister contact frequency P(s) as a function of genomic separation s for the two sisters. (F) Trans-sister scsHi-C contact map for the same window as in (D), plotted with Sister 1 along the y-axis and Sister 2 along the x-axis. Contacts above the diagonal (trans 1<2) are enriched relative to those below (trans 2<1). (G) Interpretation of trans-sister contact maps. Trans 1<2 contacts above the diagonal occur when Sister 1 is shifted relative to Sister 2 along 5′→3′ of the inherited (old) strands (1<2 shift); trans 2<1 ones below the diagonal require a shift in the opposite direction (5’→3’ along the new strands). (H) Plots of trans-sister contact frequency P(s) as a function of genomic separation s, plotted separately for two shift directions (top), and their ratio (bottom).

Recent advances in sister-chromatid-specific chromosome conformation capture (scsHi-C) (*30, 31*) and strand-specific fluorescence in situ hybridization (FISH) (*32–34*) now enable the probing of chromatids’ conformations and alignment for potential asymmetries. By applying these techniques in human cells, we reveal a previously unrecognized genome-wide shift in chromatid alignment in the 5′-3′ direction of the inherited DNA strands. Using polymer models, we propose how this global shift can arise from biased cohesion establishment by the replication forks. This discovery links the molecular asymmetry of DNA replication to higher-order chromatin architecture, raising new questions about the mechanisms of homology search during DNA repair.

## Results

### Strand-specific labeling distinguishes sister chromatids in scsHi-C

To investigate how sister chromatids are organized following DNA replication, we employed a chromosome conformation capture method that can distinguish between sister chromatids, sister-chromatid-sensitive Hi-C (scsHi-C) (Fig. 1C) (*30*). This technique labels newly synthesized DNA strands by incorporating the thymidine analog 4-thio-thymidine (4sT) during S phase. Subsequent chemical conversion of 4sT into 5-methyl-cytosine (T→5mC) introduces signature mutations that can be detected via next-generation sequencing and mapped to the plus or minus strands. These mutations allow us to split contacts into intramolecular (cis-sister) ones, where mutations are detected on the same strand, and intermolecular (trans-sister) ones, where mutations occur on different strands (*30, 31*).

To resolve the conformation of individual sister chromatids and their relative positioning, we developed an analytical framework that assigns interacting DNA fragments to one of the two sisters based on the strand containing these signature mutations (Fig. 1C, supplementary materials). DNA fragments with T→C mutations on the minus strand are assigned to Sister 1 (inheriting the parental plus strand), whereas segments with T→C on the plus strand are assigned to Sister 2 (inheriting the old minus strand). Complementary A→G mutations arising during PCR amplification are interpreted oppositely: A→G on the plus strand indicates Sister 1, and A→G on the minus strand indicates Sister 2. This strategy yields separate cis-sister contact maps and reveals the directionality of trans-sister interactions, enabling a detailed analysis of sister chromatid alignment.

To investigate whether sister chromatids differ in their loop architectures, we reanalyzed published scsHi-C data from HeLa cells (*30*) using our new sister-resolving framework. This analysis produced two separate cis-sister contact maps: one for Sister 1, which inherits the parental plus strand, and one for Sister 2, which inherits the minus strand (Fig. 1D). Each map contained approximately 9 million interactions between DNA sites separated by more than 10 kilobases. Side-by-side visualization of a representative 7-megabase region on chromosome 1 at 100-kilobase resolution revealed that the overall folding patterns, including the position and strength of topologically associating domains (TADs), were strikingly similar between the two sister chromatids.

To further quantify these similarities across the genome, we compared two key measures of loop organization. First, we examined the scaling behavior of contact probability with increasing genomic distance, known as the P(s) curve (Fig. 1E), which provides insight into average loop length and density (*7, 35*). Second, we measured contact enrichment around CTCF binding sites at 5-kilobase resolution, since these sites are known to stabilize cohesin-mediated chromatin loops (*36, 37*) (fig. S1A). Both analyses revealed highly concordant contact profiles between Sister 1 and Sister 2, with no discernible differences in either the P(s) scaling or the loop footprints surrounding CTCF sites. Together, these results demonstrate that loop architectures are preserved between sister chromatids, suggesting that chromatin folding is largely symmetrical after replication, regardless of which DNA strand is inherited.

### Trans-sister contacts reveal the asymmetric alignment of sister chromatids

To investigate potential asymmetries in sister chromatid alignment, we visualized the trans-sister contact map analogously to a trans-chromosomal map, plotting coordinates from Sister 1 on the vertical axis and Sister 2 on the horizontal axis (Fig. 1F). These maps can be divided along the diagonal to detect directional bias in trans-contacts (Fig. 1G). The upper triangle contains contacts between lower-coordinate loci on Sister 1 and higher-coordinate loci on Sister 2 (trans 1<2), indicating a shift in the 5′→3′ direction along the inherited strands. The lower triangle represents contacts of the opposite type (trans 2<1) that require a shift along the 5′→3′ direction of the new strands.

Strikingly, we observed a pronounced enrichment of 1<2 contacts (upper triangle), suggesting a systematic shift in sister chromatid alignment (Fig. 1F, fig. S2B). To quantify this bias, we computed the genome-wide frequency of trans 1<2 and 2<1 contacts as a function of genomic separation (Fig. 1H). The resulting curves showed a pronounced enrichment of trans 1<2 contacts at separations between 100 kb and 10 Mb, with a peak ∼2-fold asymmetry at ∼1 Mb, consistent across all chromatid pairs (fig. S1C). Taken together, scsHi-C reveals a systematic megabase-scale shift in chromatid alignment in the 5’→3’ direction along the inherited strands.

### Sister-chromatid-sensitive FISH confirms sister chromatid shift

To test whether scsHi-C contact asymmetry reflects a physical shift between sister chromatids, we performed sister-specific DNA imaging using RASER-FISH (Resolution After Single-strand Exonuclease Resection Fluorescence In Situ Hybridization) (*33, 34*). In this technique, newly synthesized DNA strands are labeled with UV-sensitive nucleoside analogs BrdU/C during S phase, then selectively nicked by UV irradiation and degraded by exonucleases (Fig. 2A). This treatment exposes ssDNA stretches of the inherited strands and enables sister-specific FISH using probes complementary to plus or minus DNA strands (Fig. 2A).

**Figure 2.**
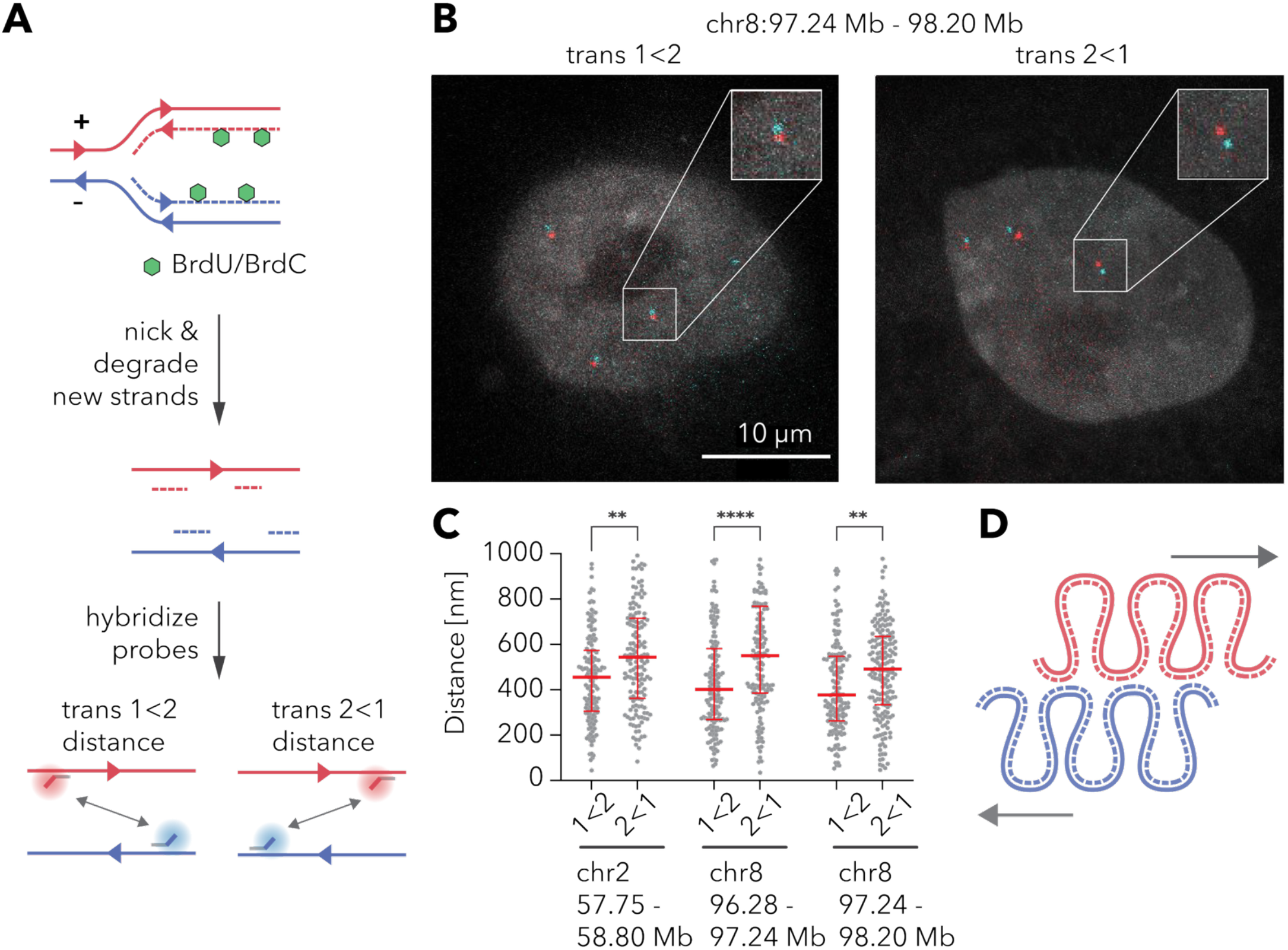
Sister-specific RASER-FISH confirms that sister chromatids are preferentially shifted 5′→3′ along the inherited strand. (A) Principle of sister-chromatid-sensitive RASER-FISH. Newly synthesized DNA is pulse-labeled with BrdU/BrdC (green). After UV nicking and exonuclease digestion of the new strands, parental strands (solid lines) remain intact, allowing strand-specific probes to hybridize. Comparing spatial distances for two different probe arrangements, 1 < 2 (bottom left) and 2 < 1 (bottom right), enables measurement of sister chromatid shift. (B) Representative mid-nuclear sections from G2 HeLa cells hybridized with probes flanking a 1 Mb interval on chromosome 8 (97.2–98.2 Mb). Red and cyan dots mark sister-specific probes. Insets zoom in on one pair of probes; scale bar, 10 μm. The image shows three pairs of dots due to a duplication of the corresponding locus in HeLa. (C) Swarm plots of center-to-center distances for three independent probe sets (∼1 Mb apart) on chromosomes 2 and 8. For every locus pair, distances are systematically shorter in the 1 < 2 orientation than in the 2 < 1 orientation. The red bars show median and interquartile range. (D) Cartoon summary of the sister chromatid shift independently measured by scsHi-C and RASER-FISH. Sisters 1 (red) and 2 (blue) are systematically shifted in the 5’→3’ direction of the inherited DNA strands.

To quantify the sister chromatid alignment shift, we measured trans-sister spatial distances between pairs of loci in two configurations: “1<2”, i.e., between a lower-coordinate site on Sister 1 and a higher-coordinate site on Sister 2, and “2<1”, the opposite (Fig. 2A, bottom). We reasoned that a systematic difference between these two measurements would indicate a shift in the chromatid alignment. We designed probe sets against three different pairs of genomic loci spaced ∼1 Mb apart - the genomic distance at which scsHi-C contact asymmetry peaks (Fig. 2B and fig. S2). In all three cases, the inter-probe distances were significantly shorter in the 1<2 configuration by 100-200 nm (Fig. 2C). Thus, RASER-FISH independently confirmed that sister chromatids are shifted in the 5′→3′ direction of the inherited strands (Fig. 2D).

### Sister chromatid shift depends on “cohesive”, but not “extrusive” cohesins

To determine whether sister chromatid shift occurs throughout entire chromosomes, we developed an asymmetry score. For each 100 kb genomic bin, this score quantifies the local asymmetry of trans-sister contacts as the log2 ratio of 1<2 and 2<1 trans-sister contacts over distances less than 1 Mb (Fig. 3A, supplementary materials). This score was consistently positive at over 97% of bins, with a genome-wide median of ∼1.0, indicating a genome-wide robust, directionally consistent shift (Fig. 3B). Although contact asymmetry was more visually apparent near TAD boundaries in scsHi-C maps (Fig. 1F and 3A), the asymmetry scores showed only minor enrichment at TAD boundaries (Fig. 3B) or regions with high trans-sister contact density (fig. S3A). The correlations with compartmentalization and replication timing were only weak (Fig. 3B and fig. S3B) (*38*), suggesting that the sister-chromatid shift is a general feature of sister chromatid organization across the genome.

**Figure 3.**
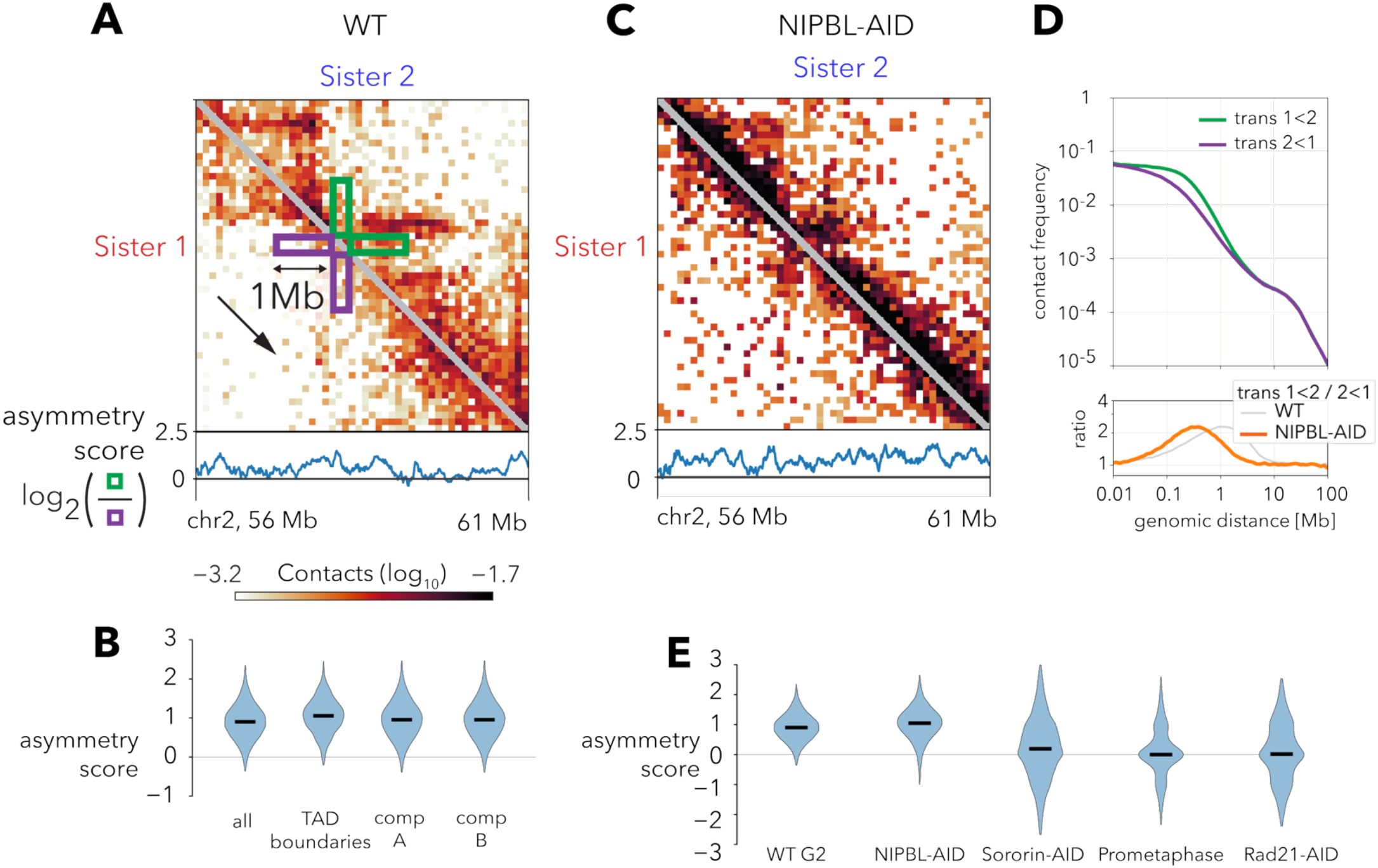
Cohesive, but not extrusive, cohesins maintain the 5′→3′ sister-chromatid shift. (A) Trans-sister scsHi-C map (100 kb bins) and asymmetry score for a 5 Mb window on chromosome 2 in wild-type (WT) G2 cells. For every position, the asymmetry score is calculated as a log2-ratio of its trans-sister contacts up to 1 Mb above and below the diagonal (the green and magenta rectangular windows, correspondingly). (B) Violin plots of the asymmetry score genome-wide (“all”), at topologically associating domain (TAD) boundaries, in A-, and B-compartment regions. Black bars show the medians. (C) Trans-sister map of the same locus as in (A) after auxin-induced degradation of NIPBL (NIPBL-AID). (D) Plots of trans-sister contact frequency P(s) as a function of genomic separation s, plotted separately for two shift directions (top), and their ratio (bottom). (E) Genome-wide asymmetry scores in WT and under conditions that selectively remove different cohesin pools: NIPBL-AID, Sororin-AID, prometaphase, and Rad21-AID. Black bars show the medians.

To test whether sister chromatid misalignment might be caused by cohesin loop extrusion, we reanalyzed published scsHi-C datasets from cells depleted of NIPBL (*39*), a cohesin subunit essential for its ATPase-driven motor activity (*3, 4, 6, 40, 41*). In these experiments, NIPBL was depleted using an auxin-inducible degron (AID) (*42*) after DNA replication, thus preserving already established cohesive links. As expected (*6*), NIPBL loss reduced known loop extrusion structures in cis contact maps (fig. S3C to E). Trans-sister contacts also became more uniform (Fig. 3C), losing the stripes of contact enrichment (fig. S3F), and concentrated along the diagonal (fig. S3G). Nevertheless, trans-sister contact asymmetry remained equally pronounced (Fig. 3D and E), with a ∼2-fold enrichment of 1<2 over 2<1 contacts at shorter distances of ∼300 kilobases. These results indicate that loop extrusion redistributes trans-sister contacts but is not essential for maintaining chromatid misalignment.

To test if cohesive cohesins maintain the sister chromatid shift, we examined three experimental conditions with impaired cohesion. First, we studied cells depleted of Sororin, a key stabilizer of cohesive cohesins, whose loss selectively removes cohesive, but not extrusive, cohesins from chromosomes (*43–45*). To this end, we reanalyzed previously published scsHi-C datasets (*30*) and performed additional experimental scsHi-C replicates. These cells showed a marked reduction in both the frequency and directional bias of trans-sister contacts (Fig. 3E and Fig. S3H). Second, we examined scsHi-C maps from prometaphase-synchronized cells, where most cohesins are removed from chromosomes by the prophase pathway (*16, 46, 47*). Consistently, these cells revealed almost complete symmetrization of trans-sister contacts (Fig. 3E and fig. S3I). Finally, we performed an AID depletion of the essential cohesin component Rad21, removing both extrusive and cohesive cohesins. In scsHi-C, these cells too showed a symmetric trans-sister contact profile (Fig. 3E and fig. S3J). Together, these findings demonstrate that the sister chromatid shift requires the presence of cohesive cohesin and is not a consequence of loop extrusion. Thus, cohesive cohesins not only tether sister chromatids together but also maintain their directional misalignment.

### Directionally misaligned cohesive cohesins can induce sister chromatid shift

The simplest explanation for sister chromatid shift is that cohesive cohesins connect the two chromatids at systematically mismatched positions (Fig. 4A). To test if such sparse misaligned links (estimated at 1 per ∼100 kb (*48*)) could shift entire chromosomes by ∼300 kb, we turned to polymer simulations. We modeled sister chromatids as flexible chains connected by cohesins and undergoing Brownian motion (Fig. 4B). To match the trans-sister P(s) from NIPBL-depleted cells (Fig. 4C), our models required two key features (fig. S4A): (1) an average misalignment of cohesins in the 1<2 direction by ∼100 kb (Fig. 4D and fig. S4A), and (2) additional symmetric misalignment by ∼100-150 kb per anchor (fig. S4A). This combination created an excess of positively misaligned links with a peak at ∼300 kb in log-space (fig. S4B) and reproduced the experimental trans-sister P(s) curves, symmetric at short distances (<30 kb), but asymmetric at intermediate scales (30 kb - 3 Mb, Fig. 4C). The best-fitting model inferred the inter-cohesin distance of ∼100 kb (Fig. 4D and fig. S4C and D), remarkably consistent with independent estimates (*48*). Thus, modest (∼100 kb) biased misalignment of cohesive links is sufficient to induce the megabase-scale chromatid shift observed experimentally.

**Figure 4.**
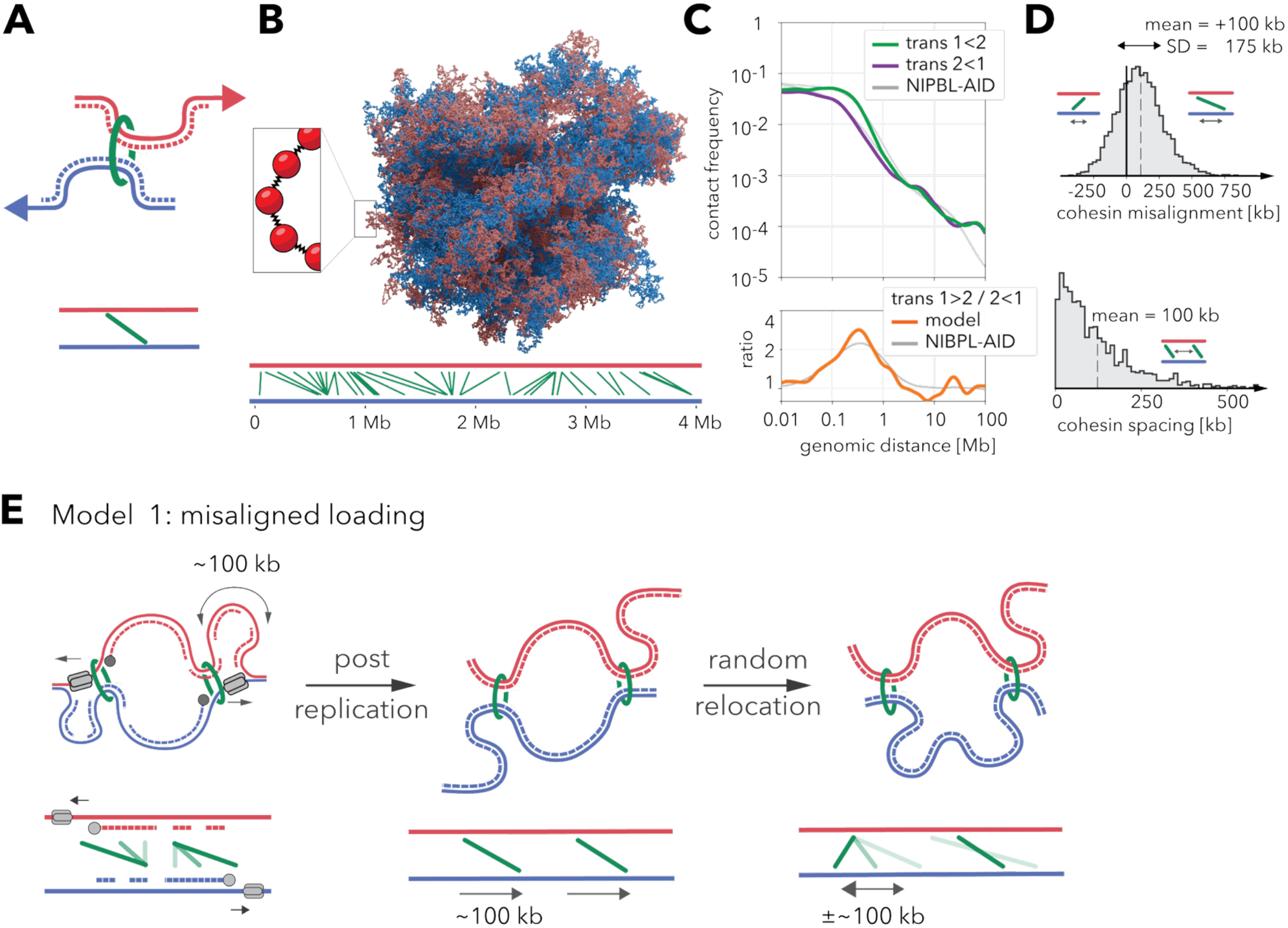
Directional misalignment of cohesive cohesins by replication forks can induce a sister chromatid shift. (A) Top: A misaligned cohesive cohesin (green), that tethers two loci on Sisters 1 and 2 (red and blue) in a 1<2 configuration, induces a local sister chromatid shift in the 5’→3’ direction of inherited strands (arrows). Bottom: a corresponding 1D diagram showing the position of sites on two sister chromatids (red and blue horizontal axes) bridged by the cohesin (green segment). (B) Top: an example 3D conformation of the best-fit model of cohesed sister chromatids. The inset illustrates that chromatin is modeled as a polymer chain of beads connected by springs. Bottom: corresponding 1D diagram of cohesive cohesin trans-sister links within a 4 Mb segment of the two modeled sisters. (C) Top: *in silico* trans 1<2 and 2<1 contact frequency P(s) curves (top) and their ratio (bottom) for the best-fitting model of sister chromatids linked by misaligned cohesins. Gray lines show the P(s) curves for the NIPBL-AID scsHi-C data (same as in Fig. 3D). (D) Histograms showing the distributions of cohesin misalignments (top) and separations (bottom) in the best-fitting model shown in (B) and (C). (E) Top: a model for the establishment of misaligned cohesion during replication. Left: cohesins move along with the replication machinery on the continuously synthesized leading strand, while remaining stationary along the discontinuously synthesized lagging strand. This difference in movement generates transient DNA loops that connect the cohesin’s original position on the lagging strand to its advancing position on the leading strand. Center: After replication, these loops resolve into stable cohesive links that connect misaligned sites. Right: Subsequent random repositioning perturbs these cohesin positions symmetrically in both directions. Bottom: corresponding 1D diagrams of cohesin positions; in the right panel, the color gradient shows the temporal progression of misalignments (from light to dark).

### Model 1: Replication forks misalign cohesins asymmetrically during loading

What molecular mechanisms can systematically misalign cohesins? Replication is the only known source of difference between sisters (*17*); replication forks establish cohesion and are inherently asymmetrical. Thus, we identified two mechanistic models of replication-coupled cohesion misalignment, both consistent with current observations.

In the first model, cohesins are directly misaligned by forks during DNA replication (Fig. 4E). Once captured by forks, cohesins travel with the replication machinery along the leading strand, while remaining stationary along the discontinuously synthesized lagging strand (Fig. 4F, left). This differential movement creates transient DNA loops that bridge the cohesin’s initial lagging strand position to its advancing leading strand position. Upon fork passage and cohesin release, these loops resolve into stable links connecting misaligned sites (Fig. 4E, middle) separated by up to half the replicon size, i.e., ∼75-100 kb (*49*). Subsequently, diffusion or random displacement by DNA-tracking motors perturb these positions symmetrically by an extra ∼100-150 kb (Fig. 4E, right; also see simulations in fig. S4E and supplementary materials), generating the inferred distribution of misalignment.

### Model 2: “paired semi-anchored cohesion” misaligns asymmetrically via random sliding

Alternatively, cohesins may be initially loaded in-register but develop asymmetry through constrained sliding. In this model, individual cohesins are “semi-anchored”, i.e., can diffuse along one sister chromatid but remain fixed to the other (Fig. 5A). When adjacent cohesins are semi-anchored to different sisters, they form constrained pairs that can slide apart but cannot pass each other, creating local asymmetry (Fig. 5B, top). The direction depends on anchoring order, e.g., “2-1” pairs (first cohesin anchored to sister 2, second to sister 1) generate 1<2 misalignment (Fig. 5B, bottom). Replication bubbles can establish “2-1” pairs across the genome by accumulating cohesins at termination sites (as suggested in (*13*)) and anchoring them preferentially to the leading strands (Fig. 5C).

**Figure 5.**
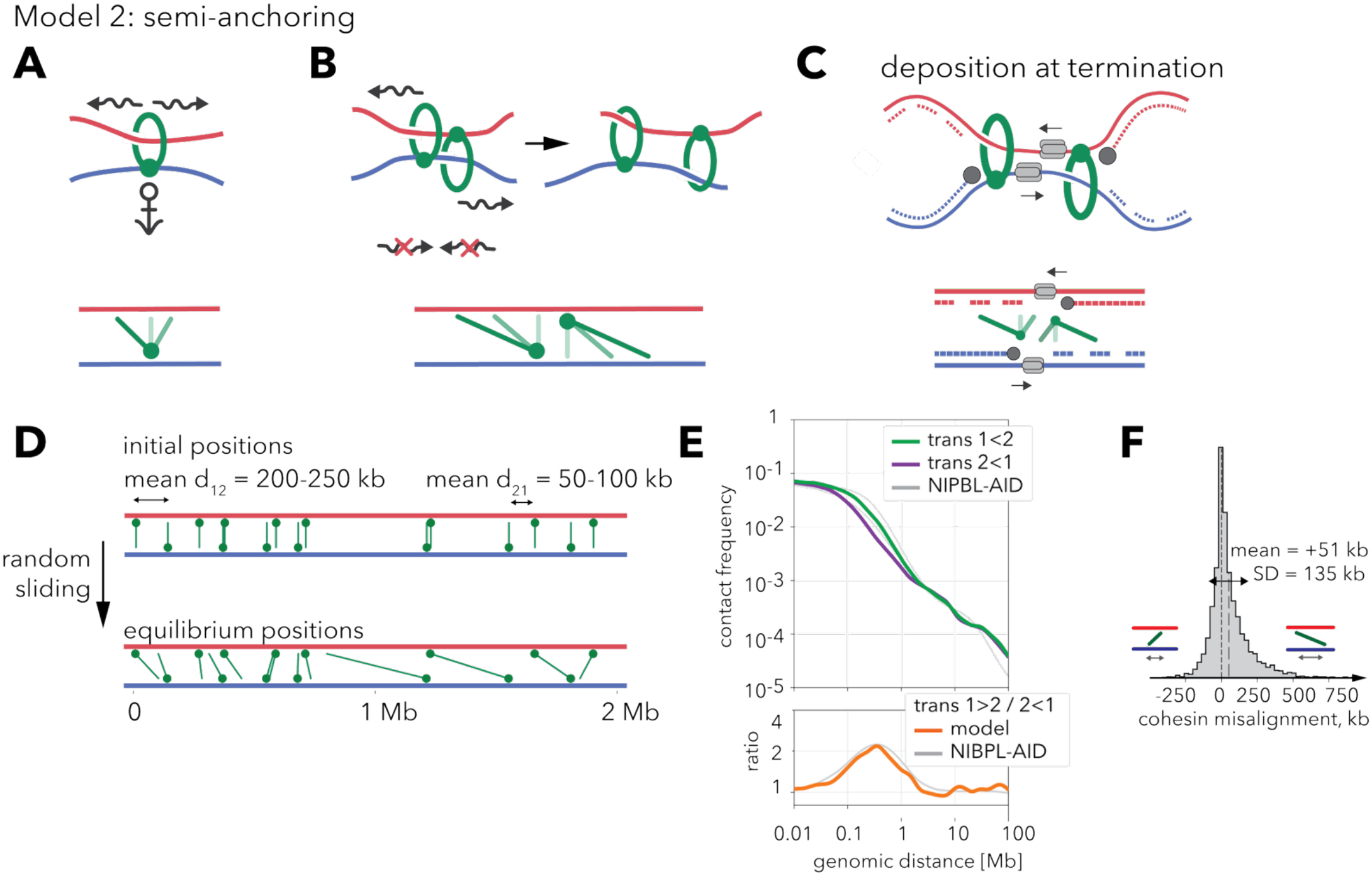
Pairs of cohesins semi-anchored to opposite sister chromatids can induce a sister chromatid shift. (A) Top: an illustration of the semi-anchored cohesin hypothesis, which postulates that cohesins are tethered to one sister chromatid but can slide along the other one. Bottom: The corresponding 1D diagram of trans-sister links; the color gradient illustrates a few permitted link orientations. (B) Top: an illustration of a pair of semi-anchored cohesins fixed to opposite sister chromatids. The rings can move apart (right), but cannot slide past each other (red crossed arrows), resulting in an asymmetry in their sliding direction (left). Bottom: same as in (A). (C) Top: an illustration of a pair of oppositely semi-anchored cohesins at converging asymmetric replication forks. Anchoring at the leading strand may be mediated by interactions with the CMG helicase (grey cylinders), polymerase ε (gray circles), or other elements of the fork. Bottom: same as in (A). (D) 1D diagram of trans-sister links in the best-fit model of semi-anchored cohesion. Top: initially, cohesins are randomly placed in alternating anchoring order, thus forming “1-2” and “2-1” pairs with different average spacings d_12_ and d_21_. Bottom: in the simulations, the free side of each link can diffuse freely up to the anchors of the two adjacent cohesins. After a sufficiently long time, the distribution of links reaches a steady state. (E) Top: *in silico* trans 1<2 and 2<1 contact frequency P(s) curves (top) and their ratio (bottom) for the best-fitting model of cohesed sister chromatids. Gray lines show the P(s) curves for the NIPBL-AID scsHi-C data (same as in Fig. 3D). (F) The equilibrium distribution of cohesin misalignments in the best-fitting model.

To test whether this mechanism can generate the observed genome-wide sister chromatid shift, we modeled semi-anchored cohesins as slip-links statically attached to one sister and freely sliding along the other (Fig. 5D and fig. S5A). Cohesins were initially loaded in-register and semi-anchored in alternating order, forming stochastically positioned “1-2” and “2-1” pairs with average spacings d_1-2_ and d_2-1_, correspondingly. This model accurately reproduced trans-sister interaction frequencies in NIPBL-depleted cells (Fig. 5E) with d_2-1_ ∼50-100kb and d_1-2_ ∼200-250kb (fig. S5B). The substantial spacing within “1-2” pairs (d_2-1_>0) is required to reproduce symmetric trans-contacts at short distances (s≲30kb) (Fig. 5F and fig. S5C), and can develop from tightly deposited pairs through slow random sliding of anchors over time (see simulations in fig. S5D and supplementary materials). This model demonstrates that asymmetric cohesion and sister chromatid shift can emerge spontaneously through mechanical constraints on random cohesin sliding, without requiring active misalignment during replication.

## Discussion

Our study uncovers previously unappreciated structural asymmetry of cohesed sister chromatids. Using strand-specific Hi-C and FISH, we reveal a genome-wide directional shift between chromatids in the 5′–3′ direction of the inherited parental strands. This discovery shows that the molecular asymmetry of the DNA replication machinery imposes a directional framework on the large-scale chromatin architecture.

Central to this asymmetry is the positioning of cohesive cohesin complexes. The shift persists in the absence of loop extrusion but is eliminated when cohesion is disrupted. Polymer simulations show that a modest (∼100 kb) directional bias in cohesin placement, combined with stochastic variability, is sufficient to reproduce the observed trans-sister contact asymmetry.

We propose two mechanistic principles by which asymmetric replication forks can bias the alignment of cohesive cohesins. In the “direct misalignment” model, cohesins are displaced differentially by fork progression -actively moved along the leading strand while impeded on the lagging strand. This model predicts the formation of transient ∼100 kb cohesin-dependent loops behind the fork. In the “semi-anchored” model, asymmetric cohesion emerges through biased constraints on cohesin mobility. This model predicts that cohesive cohesins can slide along one chromatid while being anchored to the other (similar to the anchoring proposed for extrusive cohesins (*50*)) and that forks anchor cohesins to the leading strands around termination sites (*13*). This creates asymmetric cohesin pairs that slide apart predominantly in the 1<2 direction. Importantly, both models are consistent with heterogeneous cohesin populations, where only a fraction exhibits directional misalignment (see supplementary materials and fig. S6, A to D)

Both models predict that cohesive cohesins slide along chromosomes during and after S phase by hundreds of kilobases, consistent with previous qualitative observations (*51, 52*). Our predictions are quantitatively consistent with the in vitro rate of cohesin diffusion on DNA of ∼1–2 μm^2^/s (*53, 54*), which scales up to hundreds of kilobases over 12–24 hours. Furthermore, this sliding can be accelerated by interactions with moving RNA polymerases (∼1-4 kb/min (*54–57*)) or loop extruding cohesins (∼60-120 kb/min (*58*)).

Our models predict that cohesion is infrequent and misaligned, explaining previously observed separations of hundreds of nanometers between identical sister loci (*59–62*). This imperfect cohesion raises questions about how homology search during DNA repair can identify suitable repair templates, which are normally provided by the sister locus at the same genomic position as the break site (*63*). Our models provide a quantitative framework for testing hypotheses on how cells overcome these separations, e.g., through active DNA break motion (*64, 65*) or extrusion-mediated scanning (*66, 67*).

## Acknowledgements

We thank Dmitry Mylarshchikov for help with CTCF ChIP-seq analysis. We thank the members of the Goloborodko and Gerlich labs, as well as Geoffrey Fudenberg, for their productive discussions. We thank the IMBA/IMP/GMI BioOptics and the Vienna BioCenter Next Generation Sequencing facilities for technical support, and J. Ellenberg and K.S. Beckwith for technical advice on RASER-FISH. The computational results presented were obtained using the CLIP cluster.

## Funding

Research in the laboratory of AG has been supported by the Austrian Academy of Sciences, Austrian Science Fund (FWF) grant SFB F 8804-B “Meiosis”, and the European Research Council (ERC) under the European Union’s Horizon 2020 research and innovation programme (grant agreement no. 101163751). Research in the laboratory of DWG has been supported by the Austrian Academy of Sciences, the Vienna Science and Technology Fund (WWTF; project LS19-001), and the European Research Council (ERC) under the European Union’s Horizon 2020 research and innovation programme (grant agreement no. 101019039). DWG is also an adjunct professor at the Medical University of Vienna. TLS has received fellowships from the European Union’s Framework Programme for Research and Innovation Horizon 2020 (Marie Curie Skłodowska Grant Agreement Nr. 847548), the European Molecular Biology Organization (ALTF 866-2022), and from the European Union’s Horizon Europe research and innovation programme (Marie Skłodowska-Curie Individual Fellowship 101103258 – MicroChrom). SK received a PhD fellowship from the Boehringer Ingelheim Fonds. Flavia Corsi has received funding from the Vienna International Postdoc Program (VIP2) and the European Union’s Horizon 2020 research and innovation programme under the Marie Sklodowska-Curie grant agreement No. 101033347.

## Author contributions

F.C. designed and performed polymer simulations. S.K. and A.G. analyzed and visualized scsHi-C data. T.S. designed, performed, and analyzed RASER-FISH experiments. Z.T. and P.B. performed scsHi-C experiments. M.M., D.W.G, A.G. performed the initial analyses of the scsHi-C asymmetry. F.C., D.W.G., and A.G. designed the project, analyzed data, and contributed to writing the manuscript. F.C., S.K., T.S., D.W.G., and A.G. acquired funding.

## Data and materials availability

Scripts used for data analysis and modeling will be publicly available at https://github.com/glab-vbc/sister-asymmetry-analysis and https://github.com/glab-vbc/sister-asymmetry-models.

## Notes

### Competing Interest Statement

The authors have declared no competing interest.

