## Supplementary Materials for "Conformational asymmetry of replicated human chromosomes"

### Materials and methods

#### Cell culture for scsHi-C and RASER-FISH experiments.

Cells were grown in Dulbecco's modified Eagle medium (DMEM) supplemented with 10% (v/v) fetal bovine serum (FBS) (Gibco), 1% (v/v) penicillin-streptomycin (Sigma-Aldrich, P0781 ) and 1% (v/v) GlutaMAX (Invitrogen, 35050038) supplemented with 0.5 ug/ml puromycin (Calbiochem, 540411). The cell lines were cultured in a humidified incubator at 37°C, with 5% $CO_2$.

Both HeLa RAD21-mEGFP-AID and HeLa EGFP-AID-Sororin cell lines used in this study were derived from the HeLa Kyoto cell line, which was obtained from S. Narumiya (Kyoto University, Japan) and validated using the Multiplex Human Cell Line Authentication (MCA) test. The RAD21-mEGFP-AID cell line was previously described in (*1*), and the EGFP-AID-Sororin cell line was previously described in (*2*). Both cell lines have been regularly tested negative for mycoplasma contamination.

#### scsHi-C experiments

**Cell synchronization for scsHi-C**

For scsHiC, cell synchronization was performed as described in (*2*, *3*), with cells cultured in the absence of puromycin. Briefly, 0.5 M cells were seeded into 25-$cm^2$ flasks and grown for 3 h, then supplemented with 2 mM thymidine (Sigma-Aldrich, T1895). Cells were released 16 h later by washing twice with prewarmed medium. Eight hours later, cells were supplemented with 3 μg/ml aphidicolin (Sigma-Aldrich, A0781) and 2 mM 4sT (Carbosynth, NT06341). Cells were released 16 h later by washing twice with prewarmed medium and adding medium containing 2 mM 4sT. To deplete Sororin or RAD21, indol-3-acetic-acid (auxin)(Sigma-Aldrich, I5148) was added to the cells 1h before the second release to a final concentration of 500 uM. Four hours after release, cells were supplemented with 9 μM RO-3306 (Sigma-Aldrich, SML0569). Cells were collected 20 hours later by washing twice with PBS, followed by trypsinization and resuspension in medium. RO-3306 and auxin were added to all buffers and medium used for harvesting. Cells were then spun down, washed again with PBS, followed by fixation for 4 min in 1% formaldehyde (Life Technologies, 28906). Formaldehyde was quenched by washing the cell pellet once with 20 mM TRIS-HCl pH 7.5 (Sigma-Aldrich, T6066). Cell pellets were frozen and stored at -20°C until further processing.

**scsHi-C sample preparation**

scsHi-C samples were prepared as described in (*2*) and (*3*). In brief, fixed cells were permeabilized with an ice-cold Hi-C lysis buffer, containing 10 mM TRIS-HCl pH 8.0 (Sigma-Aldrich, T6066), 10 mM NaCl (Merck, 1.06406.5000), 0.2% Nonidet P-40 substitute (Sigma-Aldrich, 74385), 1x complete EDTA-free protease inhibitor (Roche, 11836170001) for 30 min at 4 °C. The cells were then centrifuged at 2500 g for 5 minutes, and the supernatant was discarded. Next, the cells were incubated with a digestion mix (375 U DpnII (NEB, R0543) in 1× DpnII buffer (NEB, B0543S)) at 37 °C for 16 hours under rotation. Following this step, the cells were again centrifuged, and the supernatant was discarded. A fill-in mix containing 38 μM biotin-14-dATP (Thermo Fisher Scientific, 19524016), 38 μM dCTP, dGTP, and dCTP (Thermo Fisher Scientific, R0152, R0162, R0172), 50 U Klenow Polymerase (NEB, M0210), 1x NEB 2 buffer (NEB, B7002S) was added, and the cells were incubated at 37 °C for 1 hour under rotation. The cells were again centrifuged, and a ligation mix containing 1× T4 DNA ligase buffer (Thermo Fisher Scientific, EL0011), 0.1% Triton X-100 (Sigma, X100), 100 μg/ml BSA (Sigma-Aldrich, A7030), 50 U T4 DNA ligase (Thermo Fisher Scientific, EL0011) was added, and the cells were incubated at room temperature for 4 hours. The cells were centrifuged again, and gDNA was purified using the DNeasy Blood and Tissue kit. The DNA was sheared using a Covaris S2 machine (duty cycle 10%, intensity 5.0, cycles/burst 200, 25 seconds), and double size selection was performed using AMPure XP beads (Beckman Coulter, A63881). The eluted DNA was bound to Dynabeads MyOne Streptavidin C1 beads (Thermo Fischer Scientific, 65001) by incubating in biotin binding buffer (5 mM Tris-HCl pH 7.5 (Sigma-Aldrich, T6066), 0.5 mM EDTA (AppliChem, A1103,1000), 1 M NaCl (Merck, 1.06406.5000)) at room temperature for 1 hour, followed by washing and resuspension twice with Tween wash buffer (5mM TRIS-HCl ph 7.5 [Sigma-Aldrich, T6066], 0.5 mM EDTA [AppliChem, A1103,1000], 1M NaCl [Merck, 1.06406.5000], 0.05% Tween20 [Sigma-Aldrich, P1379-250ML]) then twice with water. Library preparation was performed using the NEBNext Ultra II DNA library prep kit for Illumina (NEB, E7645) according to the manufacturer's instructions. The libraries were eluted using 95% formamide (Sigma, F9037), 10 mM EDTA (AppliChem, A1103,1000) at 65 °C for 2 minutes, followed by DNA precipitation using ethanol (Sigma-Aldrich). Finally, 4sT was converted to methyl-cytosine using OsO4 (Sigma-Aldrich, 75632) in the presence of NH4Cl (Sigma-Aldrich, 326372), followed by qPCR using Q5 qPCR master mix (NEB, M0544) from the NEBUltra Ultra II DNA library prep kit for Illumina and DNA purification using AMPure XP beads (Beckman Coulter, A63881).

#### scsHi-C data processing and analysis

##### scsHi-C data processing

scsHi-C samples were processed using a custom nextflow pipeline (https://github.com/gerlichlab/scshic_pipeline) described in (*2*, *3*) with slight modifications. Briefly, NGS reads were aligned with bwa (*4*) against a modified hg19 reference containing HeLa Kyoto single-nucleotide variants (*2*). The resulting aligned sequences were used to generate .pairsam files using pairtools (*5*), which were then sorted and deduplicated. For each scsHi-C contact, the two interacting genomic segments were sister-resolved based on the presence of signature mutations. A contacting segment was assigned to Sister 1 if it aligned to the "+" strand and contained an A->G substitution (as compared to the reference genome), or if it aligned to the "-" strand and contained a T->C conversion. Conversely, a segment was assigned to Sister 2 if it aligned to the "+" stand and contained T->C conversion, or, if it aligned to the "-" segment and contained an A->G conversion. The sister-assigned reads were then split into four groups: cis-sister 1, cis-sister 2, trans-sister 1<2 (i.e., trans-sister contacts where a segment on Sister 1 has a lower coordinate), and trans-sister 2<1 (i.e. trans-sister contacts where a segment on Sister 1 has a higher coordinate). Each of these four groups of pairs was aggregated into a separate contact map, binned at multiple resolutions, and stored in the Cooler format (*6*). A separate .cool file was generated by merging all cis and trans sister contacts together, binned at multiple resolutions, and balanced, excluding the $0^{th}$ and $1^{st}$ diagonals to ignore noninformative short-range Hi-C artefacts. Bins with the log marginal read-count less than five median absolute deviations from the median log marginal read-count of all bins in the same chromosome were excluded. The resulting weights were copied into the four individual cooler files containing cis- and trans-sister contacts.

**Contact probability P(s) curves**

Contact-probability curves for cis- and trans-sister interactions were generated at base-pair resolution using pairtools. Genomic separations (s) from 1 kb to 1 Gb were partitioned into logarithmically spaced bins, 128 bins per order of magnitude. For each bin we counted (i) the number of observed cis-contacts whose separation fell within the bin and (ii) the total number of possible intra-chromosomal locus pairs at that separation. Both counts were independently smoothed in log10(s) space using a Gaussian kernel with $\sigma = 0.03$. The smoothed ratio of these two quantities yielded P(s).

**Compartment, insulating boundary, and dot calling**

We generated the A/B compartmentalization track using *cooltools call-compartments* (*7*), applied to all (i.e. non-sister resolved) contacts in the G2 dataset (*2*) at 10kb resolution. We detected boundaries using *cooltools insulation*, applied to all G2 contacts, at 10kb resolution, with a 200kb sliding window size, and selecting boundaries with strength >0.3. Finally, we detected sites of focal contact enrichment, known as "dots" or "loops", using Mustache (*8*) applied to all G2 contacts at 5kb resolution, with a p-value threshold of 0.1, and a sparsity threshold of 0.5.

**Hi-C aggregate maps at CTCF sites**

To calculate the average scsHi-C contact footprint of CTCF sites, we reprocessed the published dataset on CTCF ChIP-seq data from Kyoto cells (*9*). Briefly, the published reads were aligned against hthe g19 reference genome, and signal peaks were detected using MACS2 (*10*, *11*). The peaks were overlapped with CTCF motifs detected in hg19 using FIMO (*12*); for each peak, the motif with the lowest p-value was selected. Finally, we selected 48446 peaks with the maximal possible score of 1000.

Hi-C aggregate maps were computed using cooltools (*7*). For each set of genomic locations (CTCF peaks, loops, trans-sister stripes), we extracted snippets of balanced scsHi-C cis- or trans-sister contact maps centered on each location within the set. For trans-sister stripes, we averaged the value of each pixel across the stack of snippets. For CTCF peaks, we generated observed-over-expected scsHi-C aggregate maps by normalizing the value of each pixel in each stack ("observed") by the genome-wide average contact frequency of the corresponding diagonal ("expected"), and then averaging the stack of normalized snippets pixel-wise.

**Cross-score and trans-sister contact stripe calling.**

To detect stripes of trans-sister contact frequencies, we used cross-score, first introduced in (*13*) and available in cooltools (*7*). Briefly, the cross-score of each genomic bin is calculated as the sum of contacts formed it within a given range of genomic separations. To detect stripes of trans-sister contacts, we calculated cross-scores for G2 trans 1<2 and 2>1 interactions combined, at a resolution of 10kb and a separation range of 30 kb - 1 Mb. We then smoothed the resulting track using a Gaussian kernel with a standard deviation of 30 kb, and found all of their local maxima. Next, we clustered the resulting maxima using the Agglomerative Clustering algorithm of scikit-learn (*14*) in the space (peak cross-score value; log10(peak prominence)) with k=2 and linkage='ward'. Finally, we selected the high-prominence cluster, thus yielding a set of 4078 high-confidence trans-sister contact stripes.

##### Asymmetry score

We calculate the asymmetry score as the log2-ratio of cross scores calculated for trans 1<2 and 2<1 cross-scores separately:

$AS = \log_2 (\text{x-score}_{\text{trans } 1<2} / \text{x-score}_{\text{trans } 2<1})$

For plots of asymmetry scores in WT and NIPBL-AID shown in Fig. 3, we calculated asymmetry scores at 10kb resolution over a range of genomic separations 30kb-1Mb and smoothed the resulting tracks using a 100kb square kernel. For all violin plots shown in Fig. 3 and fig. S3, we calculated asymmetry scores at 100kb resolution, using a range of genomic separations of 100kb-1Mb and performing no smoothing.

#### Published datasets used in this study

We have used the previously published scsHi-C datasets with ENA accession numbers PRJNA639130 and PRJNA946666 and the GEO RepliSeq dataset accession GSM923449.

#### RASER-FISH

##### FISH methodology

FISH probe synthesis and hybridization were carried out as previously described (*15*), incorporating RASER-FISH (*16*) and oligo pool amplification (*17*).

##### FISH probe design and amplification

A HeLa Kyoto reference genome (hg19 with SNP correction (*2*) was used for primary FISH probe design. OligoMiner (*18*) was used to parse selected 30-40 kb-long target regions for 36–42 bp unique oligo sequences (42 °C annealing temperature (at 50% formamide concentration), 30–70% GC content, 0.17–0.90 LDA stringency, 5–16 k-mer filtering, and a 0.1 secondary structure filter). The stringency was varied to design ~300 probes per selected region, and probes were distributed in an alternating pattern across the two sister chromatids (~150 probes/sister). Unlabeled primary oligos were designed to contain short barcode sequences (unique sister-specific barcodes and a mutual region-specific barcode) that enable binding to fluorescently labeled secondary oligos, as well as flanking primers incorporating the T7 in vitro transcription (IVT) promoter for oligo pool amplification. The oligo pool was ordered from GenScript (GenTitan™ Oligo Pools). FISH probe amplification was performed by qPCR with 15 ng of the oligo pool, 2× Phusion High-Fidelity PCR Master Mix (Thermo Fisher Scientific, F531), 0.5 µM forward and reverse primers, and 1× EvaGreen (Biotium, 31000). After an initial denaturation at 98 °C for 3 min, cycles of 98 °C for 10 s, 66 °C for 10 s, and 72 °C for 15 s were repeated until the reaction approached a plateau. The amplified products were purified using the DNA Clean & Concentrator-25 kit (Zymo Research, D4033) and eluted in 50 µl ultrapure water. For IVT, 1.5 µg of the purified PCR product was incubated overnight at 37 °C using the HiScribe T7 Quick High Yield RNA Synthesis Kit (NEB, E2050) with 80 µl of NTP buffer mix, 6.25 µl T7 RNA polymerase mix, 6.25 µl RNAsin Plus RNase inhibitor (Promega, N2615), and ultrapure water to reach a final volume of 160 µl. Subsequently, 150 µl of the resulting ssRNA was used directly for reverse transcription (RT) at 50 °C for 1 h in a reaction containing 21 µl of 25 mM dNTP mix (final concentration 1.75 mM), 57 µl of 100 µM forward primer (final concentration 19 µM), 6 µl RNAsin Plus RNase inhibitor, 6 µl Maxima H Minus Reverse

Transcriptase (1,200 U, Thermo Fisher Scientific, EP0753), and 60 µl of 5× RT buffer. RNA was then degraded by adding 150 µl of 0.5 M EDTA and 150 µl of 1 M NaOH, followed by incubation at 95 °C for 15 min. The resulting ssDNA oligo pool was purified with a DNA Clean & Concentrator-100 kit (Zymo Research, D4030), eluted in 200 µl ultrapure water, and quantified by Nanodrop. Fluorescently labeled secondary oligos were generated via click chemistry (ClickTech Oligo Link Kit, baseclick) with 3′-C3-azide oligos (Metabion) and an alkyne-modified ATTO565/ATTO647N/ATTO488 fluorophore (ATTO-TEC, AD 488/AD 565/AD 647N). The labeled oligos were purified with two n-butanol washes (Sigma-Aldrich, B7906) and diluted in 1× TE buffer.

**Cell synchronization to the G2-phase and RASER-FISH protocol**

HeLa cells were seeded in Ibidi µ-Slide VI 0.5 Glass Bottom slides at 2.5×105 cells/ml (120 µL per channel) in the presence of 2mM thymidine (Sigma-Aldrich, T1895) to synchronize them at the G1/S boundary. Cells were released after ~18 h through 4x washes with prewarmed WT medium. Eight hours later, cells were supplemented with 3 µg/mL aphidicolin (Sigma-Aldrich, A0781) and 40 µM BrdU/BrdC (3:1 ratio; Sigma-Aldrich, B5002; Santa Cruz Biotechnology, sc-284555) to synchronize again at the G1/S boundary. Cells were released 16 h later through 4x washes with prewarmed WT medium supplemented with 40 µM BrdU/BrdC. 4 h after release, cells were additionally supplemented with 30 µM RO-3306 (Sigma-Aldrich, SML0569) for 16 h for synchronization to G2. Cells were fixed at room temperature (RT) in 4% paraformaldehyde (Electron Microscopy Sciences, 15710) in PBS for 15 min, followed by quenching with 100 mM NH4Cl (Sigma-Aldrich, 326372) for 10 min. Permeabilization was then carried out at RT for 20 min in PBS containing 0.2% Triton X-100 (Sigma-Aldrich, 327371000). After a 15 min incubation with 0.5 µg/ml DAPI in PBS at RT, cells were UV-irradiated in a Vilber Bio-Link crosslinker (254 nm, 3.6 J/cm2) to nick BrdU/C-labeled strands, which were subsequently digested with Exonuclease III (1 U/µl; NEB, M0206) at 37 °C for 15 min. Cells were then pre-incubated for 1 h at 37 °C in FISH hybridization buffer (50% (v/v) formamide (Thermo Fisher Scientific, AM9342), 10% (w/v) dextran sulfate (Merck, S4030), 2×SSC (Thermo Fisher Scientific, AM9763)), followed by an overnight incubation at 42 °C in the same buffer containing primary FISH probes (~2 nM/probe). Two washes in 50% (v/v) formamide/2×SSC at 35 °C (5 min each) were performed, followed by a wash in 2×SSC + 0.2% (v/v) Tween-20 (Sigma-Aldrich, P9416). RNA-DNA hybrids were removed via RNase H (0.05 U/µl; NEB, M0297) treatment for 20 min at 37 °C, samples were subsequently washed 2–3× in 2×SSC + 0.2% Tween-20 at RT. Secondary FISH hybridization was performed at RT for 3 min in a buffer containing 5% (w/v) ethylene carbonate (Sigma-Aldrich, E26258) and 20 nM fluorescent secondary oligos in 2×SSC + 0.2% Tween-20. Excess secondary oligos were removed by a 1-min wash at RT with 10% (v/v) formamide in 2×SSC + 0.2% Tween-20. Finally, cells were transferred to 2×SSC containing 0.2 µg/ml DAPI before imaging. All FISH experiments were visualized on a Zeiss LSM980 confocal system equipped with a 63× NA 1.4 oil DIC Plan-Apochromat objective (ZEN Blue 2020 software v3.7), in confocal mode using a z-sectioning of 200 nm.

**RASER-FISH – Image analysis**

Image analysis was performed in Fiji (Version 2.14.0/1.54f) (*19*). All multi-color FISH images were registered to correct for chromatic aberrations as previously described (*20*). For this purpose, an image stack of immobilized TetraSpeck™ Microspheres, 0.2 µm (Thermo Fisher Scientific, T7280, prepared in-house) was acquired using the same image acquisition parameters as for the FISH images, and a custom-written Fiji Macro was used to correct the offset between the color channels. Afterwards, a maximum projection of the FISH channels was generated and the distance of the correct sister-specific FISH pairs – based on the signal from the regional barcode (ATTO488) – measured using the ComDet plugin (v0.5.5 (oval ROI shape, particle size: 5 pixels, intensity threshold: 6-8 SD)) (https://github.com/UU-cellbiology/ComDet). From the obtained X,Y coordinates of the FISH pairs, the 2D Euclidean distances were calculated across sister chromatids.

#### Computational models of cohesed sister chromatids

**3D polymer model of cohesed chromatids.**

To provide structural and mechanistic interpretation to our scsHi-C data obtained on G2 chromosomes in extrusion-suppressed NIPBL-AID cells, we developed polymer models of cohesed sister chromatids in the absence of loop extrusion. The simulations were implemented in HOOMD-blue, a GPU-accelerated molecular dynamics package v3.10.0 (*21*), similarly to as previously described (*22*).

Each sister chromatid was modeled as a chain of 500,000 particles, representing a 10 nm nucleosome (200 bp of DNA), simulating 100 Mb per chromatid. In HOOMD's normalized unit system, particle mass and diameter were set to 1.0, where 1 length unit equals 10 nm in real coordinates. We used the following forces:

- Excluded volume: all particles repelled each other with Dissipative Particle Dynamics (DPD) repulsion potential ($A_{rep}$ = 10.0 $k_BT$). This force prevented spatial overlap while still allowing some strand passing for efficient topological equilibration.
- Chain connections: Consecutive particles on each chromatid were linked via harmonic bonds (equilibrium length: 1 length unit, stiffness: 100 $k_BT$/[length unit]^2), ensuring maximum bond extension below 2 length units.
- Cohesion links: Inter-sister chromatids were cohesed using harmonic bonds (equilibrium length: 1 length unit, stiffness: 0.6 $k_BT$/[length unit]^2) with an average bond length of ~4-5 length unit (~40-50 nm, mimicking cohesin size) and maximum bond length <6 length unit. In all simulations, cohesion links acted as barriers to one another, preventing mutual crossing during diffusion along the chains.

Chromosomes were confined by periodic boundary conditions to the monomer density of 0.1 per [length]^3. Simulations were initialized with two parallel chromatids linked by cohesion bonds and equilibrated using a Dissipative Particle Dynamics (DPD) thermostat (target temperature T=1 [temperature], friction coefficient $\gamma$=10.0 [mass/time]). Smaller $A_{rep}$, $\gamma$, and cohesion link bond lengths yielded similar results to the ones reported.

Each simulation was executed on one NVIDIA GPU (P100, V100, RTX 6000, A100) on the CLIP cluster at Vienna Biocenter. We ran each simulation for at least 1e7 steps (dt = 0.05 [time]) to ensure thermal equilibration, as indicated by the convergence of contact probability scaling curves P(s). Data were analyzed in Python and visualized with Matplotlib (*23*). Specific details regarding the distribution of cohesion links, simulation design, the exact number of replicates, and time steps are provided below for each type of simulation.

**Model 1: "direct misalignment" of cohesive links by the replication forks.**

The folding of sister chromatids in the 3D model above is critically controlled by the positions of cohesin links, each defined by the pair of particle indices on the two chains that it links together ($x^1_i$, $x^2_i$). Each link is described by its mean position ($x^c_i = x^2_i - x^1_i$ ) and misalignment ($m_i = x^2_i - x^1_i$, positive values indicate misalignment in the 1<2 orientation, negative - in the 2<1 orientation). To generate randomly positioned cohesin links with the specified density and distribution of misalignments, we developed the following procedure:

1) Calculate the required number of cohesins as $N_{coh} = L / d$, where L is the length of each chromatid and d is the specified cohesin density [cohesins / kb].

2) Draw $N_{coh}$ random misalignments $m_i$ from the specified distribution of misalignments (see below). Positive numbers indicate the misalignment of a given cohesin in the 1<2 orientation, negative - in the 2<1 orientation.

3) Sort links in the order of their increasing misalignment, from negative to positive, and place them onto the two chains sequentially, one after the other ($x^1_{i+1} = x^1_i + 1$). This step is guaranteed to generate an arrangement of links, where no pair crosses each other.

4) Shuffle bonds iteratively, selecting one bond at a time and moving it to a new, random position, ensuring it does not cross any existing bonds. Repeat until the convergence of the statistical properties of generated bond distributions (10*$N_{coh}$ times).

As a result, this procedure generates a collection of randomly positioned cohesin links, with a predefined distribution of misalignments. We tested several stochastic distributions of cohesin misalignments:

- Exponential distribution, with 1 parameter - the mean $\mu^e$. This distribution is strictly positive and thus inherently imposes a one-directional (asymmetric) misalignment on cohesin links. Having systematically explored the simulations with different values of $\mu^e$, we have found that it can produce the gap between 1<2 and 2<1 trans-sister P(s), but it fails to match experimental P(s), that contain almost no gap at shorter genomic separations ($s \lesssim 30$ kb) (fig. S4A).
- Gaussian distribution, with 2 parameters - the mean $\mu^g$, and standard deviation $\sigma$. When $\mu^g=0$, this distribution is symmetric around zero, producing no directional bias in cohesin alignments and symmetric trans-sister contact statistics in 1<2 and 2>1 directions. In contrast, a Gaussian distribution with a non-zero mean $\mu^g>0$ indeed can match experimental trans-sister P(s) (fig. S4A).
- ExGaussian distribution (a sum of a Gaussian- and an exponential-distributed random number) has three parameters - the mean of the exponential term $\mu^e$, and the mean and the standard deviations of the Gaussian term, $\mu^g$ and $\sigma$. We found that, when $\mu^g$ is fixed at 0, the remaining two-parameter distribution can match experimental P(s) equally well or better than the Gaussian distribution (fig. S4A). We preferred this distribution over the Gaussian due to its better interpretability - the two components can represent two different biological mechanisms, e.g., unidirectional misalignment by replication (captured by the exponential component) and

subsequent random symmetric relocation (modeled by the zero-mean Gaussian), each contributing to the variance of the resulting distribution.

For each given 1D distribution of misaligned cohesion bonds along the chains, a 3D polymer simulation was run for 1e7 timesteps to achieve thermal equilibration. For each parameter set, 3 independent replicates were performed, and the resulting final 3D conformations were averaged to compute mean contact frequencies. The best agreement with the experimental NIPBL-AID data was achieved using an ExGaussian distribution of cohesive bonds, characterized by the parameter set: exponential mean $\mu^e$=100 kb, Gaussian mean $\mu^g$=0, and Gaussian standard deviation σ=150 kb (fig. S4C).

**Modification of model 1: strictly asymmetric initial misalignment followed by explicitly modelled diffusion.**

Next, we wanted to confirm that the Gaussian component of the exGaussian cohesin misalignment inferred by Model 1 could arise from cohesin diffusion along the chains. To this end, we designed a hybrid 3D-1D dynamics simulation (fig. S4E). Initially, cohesive links were randomly misaligned and positioned as described above, with misalignments drawn from an exponential distribution with a mean $\mu^e$ = 100 kb. To simulate the temporal diffusion of these links, we coupled 3D polymer molecular dynamics with a 1D Monte Carlo process that iteratively updates bond positions, as follows:

- 1D bond relocation MCMC step: At each iteration, a random misalignment adjustment was proposed for each cohesion bond, with a step size uniformly sampled within a range of [–5, 5] monomers (equivalent to [–1, 1] kb), applied to each bond side independently, ensuring that diffusion steps remained within the intended resolution. Steps resulting in bond crossing were immediately rejected. The energy of the remaining steps before and after the proposed move was calculated as:

$$E_{old} = \frac{K}{2}(d_{old} - d_0)^2$$
$$E_{new} = \frac{K}{2}(d_{new} - d_0)^2$$

  where $d_{old}$ and $d_{new}$ are the 3D distances between linked particles before and after the move, respectively, $d_0$ is the equilibrium bond length and $k$ is the bond force constant.
  Each proposed step was accepted if it led to a decrease in energy, or with a probability given by the Metropolis criterion:

$$P_{\text{accept}} = \begin{cases} 1, & \text{if } E_{\text{new}} \le E_{\text{old}} \\ \exp(-(E_{\text{new}} - E_{\text{old}})), & \text{otherwise} \end{cases}$$

  This ensured detailed balance and allowed the system to escape local minima by occasionally accepting energetically unfavorable moves. Random numbers for acceptance testing were drawn from a uniform distribution on [0, 1]. This process was repeated 1e3 times per bond per MCMC step.
- 3D polymer simulation step: "cohesive" harmonic bonds in the polymer model were updated accordingly, and the simulation evolved via DPD molecular dynamics for 1e3 timesteps

This two-step procedure was iterated 2e4 times, yielding a total simulation time of 2e7 3D timesteps per run. Three independent replicates were performed, and for each replicate, one conformation was saved every 2e5 timesteps (i.e., every 200 1D MCMC-3D cycles), resulting in 100 conformations per replicate. Average contact frequencies were then computed across the replicates for each of the 100 stored time points. The best agreement with the experimental NIPBL-AID data was achieved at 1.04e7 timesteps (frame number 52, fig. S4E) with a root mean square deviation (RMSD) per cohesion bond of 191kb. At this point, the average global misalignment was 96 kb, with average positive and negative misalignments of 164 kb and –88 kb, respectively.

**Model 2: oppositely semi-anchored cohesive cohesins.**

The alternative model assumes that asymmetric replication forks establish cohesion by "semi-anchoring" individual cohesins preferentially to one sister. In this model, cohesive links act as slip-links, where one end remains fixed to a specific chain position while the other end can slide along the opposite chain between neighboring anchor points (fig. S5A).

To generate randomly positioned semi-anchored cohesin links with alternating anchoring orientations, we used the following procedure:

1. Calculate the required number of cohesins as Ncoh = L / d, where L is the length of each chromatid and $d=(d_{12}+d_{21}) / 2$ is the specified average cohesin density, and $d_{12}$ and $d_{21}$ are target mean spacings within “1-2” and “2-1” cohesin pairs (see below).
2. Sequentially place cohesins onto chains. Cohesins are “anchored” to sisters in alternating order: cohesin 1 on sister 1, cohesin 2 on sister 2, cohesin 3 on sister 1, etc. The spacing between consecutive cohesins is drawn from two exponential distributions with different means, based on the orientation of their anchors - $d_{12}$ for transitions for spacings between 1-anchored and 2-anchored cohesin and $d_{21}$ for spacings between 2-anchored and 1-anchored ones. Initially, cohesins are well aligned: $x^1_i=x^2_i$.
3. Equilibrate the positions of freely sliding cohesin ends through constrained 1D diffusion between adjacent anchors. To this end, we designed a hybrid 3D-1D MCMC simulation approach, similar to that in section “Modification of model 1: strictly asymmetric initial misalignment followed by explicitly modelled diffusion.” The simulation proceeded through two alternating steps (fig. S5A):

    - 1D Monte Carlo step: displace the free end of each cohesion bond by a random step, uniformly sampled between 1 and the maximum distance to neighboring anchors. To ensure adequate sampling, this process was repeated the number of times that is equal to 20% of the maximum inter-anchor separation.
    - 3D polymer simulation step: “cohesive” harmonic bonds in the polymer model were updated accordingly, and the simulation evolved via DPD molecular dynamics for $10^3$ timesteps.

This two-step procedure was iterated $10^4$ times, yielding a total simulation time of $10^7$ timesteps per run. For each parameter combination, 5 independent replicates were performed, and the contact frequencies were calculated for the resulting conformations and averaged.

We systematically explored parameter combinations of $d_{12}$ and $d_{21}$. When $d_{12} < d_{21}$, consecutive anchors form asymmetric "1-2 pairs" that preferentially slide in the 1<2 direction due to spatial constraints. When $d_{12} > d_{21}$, the opposite bias emerges. Equal spacing ($d_{12} = d_{21}$) produces symmetric distributions. The best agreement with the experimental NIPBL-AID data was achieved using $d_{12}$ = 200 kb and $d_{21}$ = 75 kb (fig. S5B).

**Modification of model 2: anchor diffusion**

To test whether the asymmetric anchor spacing inferred by Model 2 could arise from initially tight cohesin pairs that subsequently spread through anchor diffusion, we modified the semi-anchored cohesin model. The modified procedure differed from the standard Model 2 protocol in two key steps (fig. S5D):

- Initial tight pairing: Cohesins were placed sequentially with $d_{21} = 0$ (creating tight "2-1" pairs) and $d_{12}>0$.
- Static anchor relocation: Before simulation, each anchor position was displaced by a random amount sampled from a zero-mean Gaussian distribution with standard deviation $\sigma_a$. These new anchor positions remained fixed throughout the subsequent simulation.

The remaining steps (equilibration of free ends through coupled 1D Monte Carlo and 3D molecular dynamics) followed the protocol described in the previous Model 2 section. Thus, this approach modeled two distinct types of cohesin mobility: slow diffusion of cohesin anchors along one sister and fast sliding along the other.

We systematically explored parameter combinations of $d_{21}$ and $\sigma_a$. For each parameter combination, 3 independent replicates were performed, and contact frequencies were calculated for the resulting conformations and averaged. The best agreement with the experimental NIPBL-AID data was achieved using $d_{12}$ = 150 kb, $d_{21}$ = 0 kb and $\sigma_a$ = 200 kb (Fig. S5D).

**Mixture model 1: heterogeneous cohesin population.**

To account for potential biological heterogeneity (e.g., arising from two cohesin deposition pathways), we tested whether mixtures of distinct cohesin subpopulations - one with asymmetric misalignment bias and one with symmetric positioning - could reproduce the experimental scsHi-C data.

We evaluated two mixture combinations:

1. ExGaussian + symmetric Gaussian: Combining directionally biased cohesins (exGaussian distribution with $\mu^e > 0$, $\mu^g = 0$, $\sigma>0$) with symmetrically positioned cohesins (Gaussian distribution with $\mu^g = 0$, $\sigma>0$) (fig. S6A).
2. Asymmetric Gaussian + symmetric Gaussian: Combining directionally biased cohesins (Gaussian distribution with $\mu^g > 0$, $\sigma>0$) with symmetrically positioned cohesins (Gaussian distribution with $\mu^g = 0$, $\sigma>0$) (fig. S6B).

For both mixture types, the standard deviation σ of all Gaussian components was held constant across subpopulations, reflecting shared stochastic relocation processes despite different initial alignment biases. We tested mixture ratios of 25%-75%, 50%-50%, and 75%-25% between asymmetric and symmetric subpopulations, respectively, across multiple inter-cohesin distances.

For each mixture model, cohesin links were generated by sampling the specified fraction from each distribution and randomly intermixing their positions along the chains using the bond placement algorithm described in Model 1. The subsequent 3D polymer simulation protocol (1e7 timesteps for thermal equilibration, 3 independent replicates per parameter set) followed the standard Model 1 procedure.

The best agreement with the experimental NIPBL-AID data was achieved using a 50%-50% mixture of ExGaussian + symmetric Gaussian, $\mu^e$ = 250 kb, $\mu^g$ = 0, and σ = 125 kb (Fig. S6A).

**Modification of Model 2: consecutive semi-anchored cohesins with the same-chain anchoring**
To test whether replication forks can deposit multiple cohesins per replication bubble, we extended Model 2 to allow consecutive cohesins to be anchored on the same DNA chain. In this variant, each fork deposits cohesins consistently on its respective strand, creating groups of same-chain-anchored cohesins separated by transitions to the opposite chain (fig. S6C). We implemented this model using groups of four consecutive cohesins with the following organization:

- Group structure: Two cohesins anchored on chain 2, followed by two cohesins anchored on chain 1 (pattern: 2-2-1-1).
- Spacing within groups: The spacings between the four cohesins within each group are sampled from the same exponential distribution with mean $d_{21}$.
- Spacing between groups: The distance from the last cohesin of one group to the first cohesin of the next group is sampled from an exponential distribution with mean $d_{12}$.

This arrangement creates embedded "2-1" cohesin pairs within each group while maintaining the $d_{12}>d_{21}$ parameter regime that provided optimal fit in standard Model 2. The resulting pattern could represent cohesins deposited by pairs of converging forks, each depositing cohesins with a fixed spacing.

The subsequent simulation protocol (1D Monte Carlo equilibration of free ends coupled with 3D molecular dynamics) followed the procedure described in the standard Model 2 section. For each parameter combination, 1 simulation replicate was performed The best agreement with the experimental NIPBL-AID data was achieved with $d_{12}$ = 250 kb and $d_{21}$= 100 kb (Fig. S6C).

**Mixture model 2: heterogeneous cohesin population.**
To test whether experimental scsHi-C data is consistent with coexisting cohesin populations with different mobility constraints, we extended Model 2 to include two distinct cohesin types: oppositely semi-anchored pairs and freely diffusing bonds (fig. S6D). The mixed population model was implemented as follows:

- Semi-anchored pairs: Distributed using the standard Model 2 protocol with inter-anchor distances sampled from exponential distributions (means $d_{12}$ and $d_{21}$) in the optimal parameter regime ($d_{12}$ > $d_{21}$).
- Freely diffusing cohesins: Additional cohesin bonds with both ends able to slide along their respective chains, randomly interspersed between semi-anchored pairs. These bonds were initially

placed in perfect alignment ($x^1_i = x^2_i$) and allowed to diffuse freely, but without intersecting other bonds.

- We tested mixture ratios of 25%-75%, 50%-50%, and 75%-25% between semi-anchored and freely diffusing populations, respectively, across multiple parameter combinations of $d_{12}$ and $d_{21}$.

The rest of the simulation protocol (coupled 1D Monte Carlo and 3D molecular dynamics for equilibration of mobile bond ends) was the same as in the standard Model 2 section. For each parameter and mixture combination, 1 simulation replicate was performed. The best agreement with the experimental NIPBL-AID data was achieved with a mixture of 75%-25% of semi-anchored and freely diffusing cohesins, respectively, $d_{12}$ = 250 kb and $d_{21}$= 100 kb (Fig. S6D).

**Computation of in silico P(s) curves**

To enable direct comparison with experimental scsHi-C data, we computed four P(s) curves from each polymer simulation: two cis-sister curves (representing contacts within individual chains) and two trans-sister curves (representing inter-chain contacts). The trans-sister curves were directionally resolved: the 1<2 curve was based on contacts where the linear genomic coordinate on chain 1 was smaller than that on chain 2 ($x_1 < x_2$), while the 2<1 curve captured the reverse orientation ($x_1 > x_2$). Contact detection was performed using a variable-radius approach adapted from (*22*), which accounts for potential variability in the contact radius:

- Position perturbation: displace each particle by random noise sampled from a multivariate Gaussian distribution with standard deviation $\sigma_c$ = 3.5 length units (equivalent to 35 nm), optimized to align simulated and experimental P(s) curves at the 2-3 kb scale where no dominant structural effects are expected.
- Contact detection: find all particle pairs within a 3D distance of 1.0 length using the Scipy KD-tree algorithm.
- Averaging: repeat perturbation and contact detection 10 times and average the resulting contact frequencies.

Final P(s) curves were calculated from these averaged contact frequencies using the same binning and smoothing procedure applied to experimental Hi-C pairs.

**Model selection based on comparison of simulated and experimental P(s) curves**

To determine which simulations best recapitulate the experimental scsHi-C data from NIPBL-depleted cells, we systematically explored combinations of the model parameters described above. Depending on the simulation type and the associated level of noise, we performed between one and five replicates per parameter set. For each set, we averaged the resulting four P(s) curves—two *cis* and two *trans*-sister—across replicates. All experimental and *in-silico* P(s) curves were smoothed using the procedure implemented in *cooltools* with sigma_log10=0.1 and normalized to 1.0 at a genomic distance of 2 kb. To quantify model accuracy, we calculated the root mean square deviation (r.m.s.d.) between the experimental and simulated *trans*-sister P(s) curves, for both *1→2* and *2→1* P(s), on a log10-log10 scale, across genomic distances from 10 kb to 1 Mb, distances showing most of the asymmetry we want to characterize. The larger of the two r.m.s.d. values was used as the overall mismatch score between the simulation and the experimental data.

To identify the optimal parameter combinations for reproducing experimental scsHi-C data from NIPBL-depleted cells, we systematically explored the parameter space for each model variant described above. For each parameter combination, the four P(s) curves (two cis-sister and two trans-sister) were averaged across simulation replicates. Both experimental and simulated P(s) curves were smoothed in the log-log space with a Gaussian kernel (sigma_log10 = 0.1) using the standard cooltools functionality and normalized, so that cis P(s) was equal to 1.0 at a genomic distance of 2 kb.

Model accuracy was assessed by calculating the root mean square deviation (r.m.s.d.) between experimental and simulated trans-sister P(s) curves in the log10-log10 scale. The comparison focused on genomic distances from 10 kb to 1 Mb, the range where sister chromatid asymmetry is most pronounced and biologically meaningful. Separate r.m.s.d. values were computed for both directional trans-sister curves (1<2 and 2<1), and the larger of the two values was used as the overall mismatch score to ensure that models accurately captured asymmetry in both directions. Parameter combinations yielding the lowest overall mismatch scores were identified as optimal for each model type and used for subsequent analysis and biological interpretation.

### Supplementary Figures

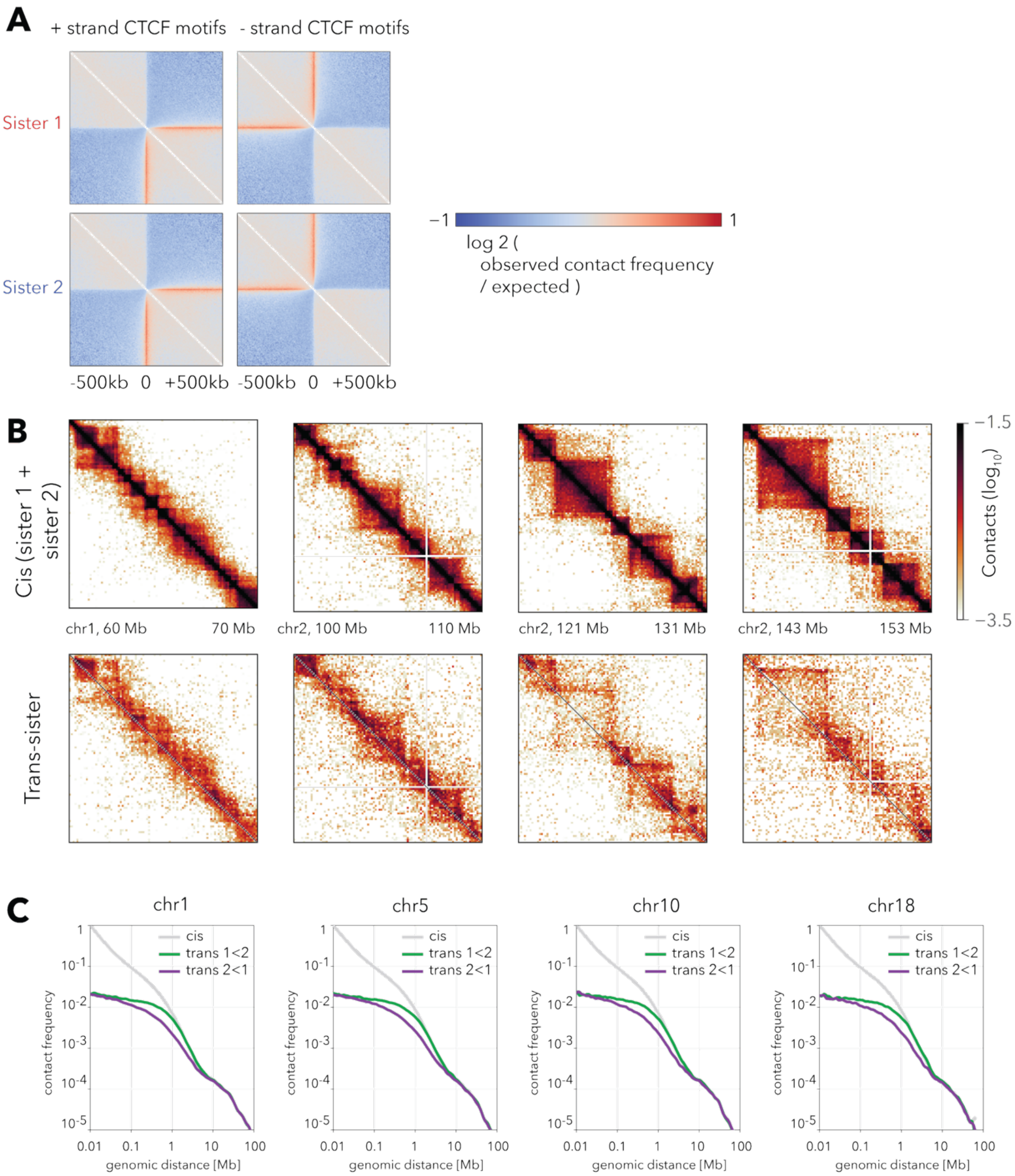


**Figure S1. Cis- and trans-sister scsH-C contact features are consistent across loci and individual chromosomes.** (A) Aggregate cis-sister contact maps (±500 kb, log2 contact frequency normalized for distance-dependent decay) centered on CTCF motifs on the + (left) or – (right) DNA strands on Sister 1 (top) or 2 (bottom). (B) Examples of cis- and trans-sister scsHi-C contact maps for a few representative 10 Mb segments of chromosomes 1 and 2, plotted at 100 kb resolution. Top row: cis-sister maps, combined for Sister 1 and Sister 2. Bottom row: the corresponding trans-sister maps. (C) Plots of cis- and trans-sister contact frequency P(s) as a function of genomic separation s for four representative autosomes (chr 1, 5, 10, 18).

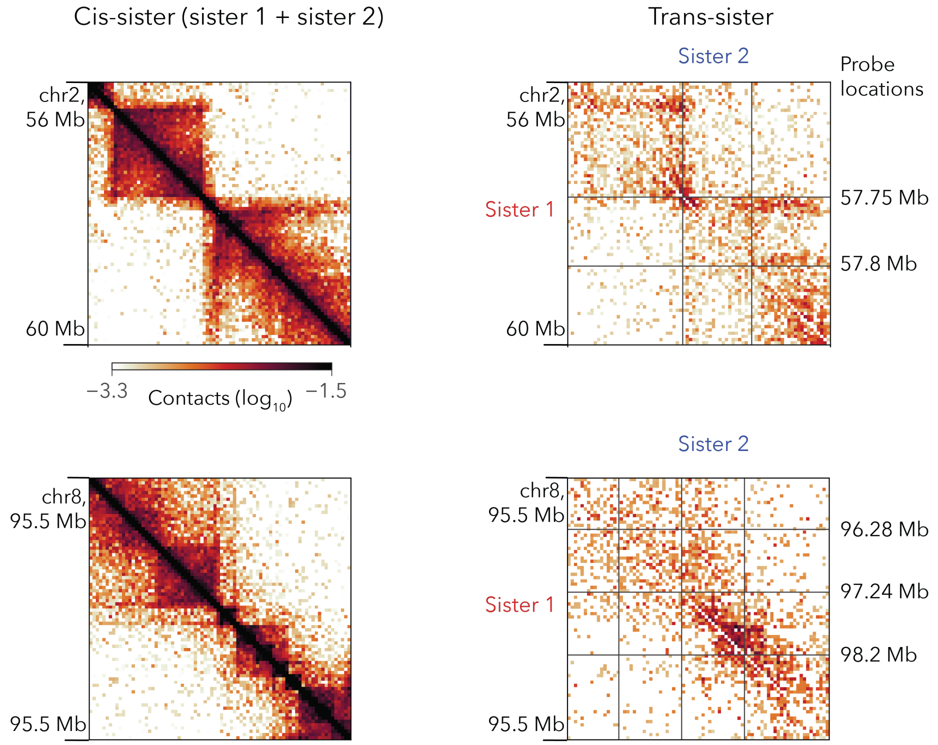


**Figure S2. The locations of the RASER-FISH probe sets.** Cis- and trans-sister scsHi-C contact maps with tagged positions of five RASER-FISH probe sets used in the study.

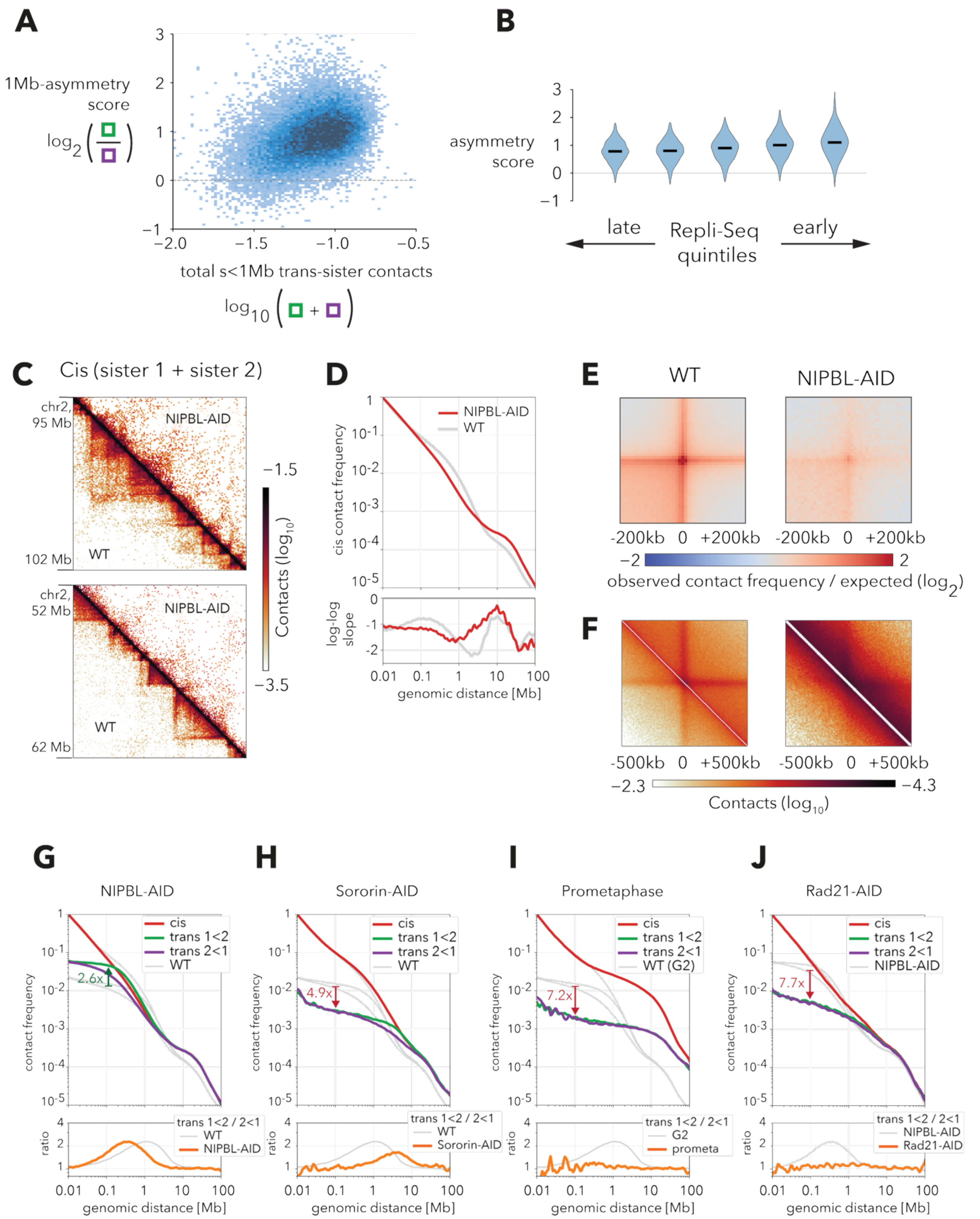


**Figure S3. Cohesive, but not extrusive, cohesins are responsible for the sister chromatid shift.** (A) A 2D histogram of scsHi-C 1Mb contact asymmetry score (y-axis) vs the total number of trans-sister <1 Mb contacts formed by each locus (x-axis). Asymmetry only weakly depends on the total number of trans-contacts and is positive for most observed loci. (B) Violin plots of the asymmetry scores of 100kb genomic bins, grouped into five quintiles based on their Repli-Seq signal. (C) The comparison of the cis-sister scsHiC contact maps in WT (lower triangle) and NIPBL-AID (upper triangle) cells in two representative loci. (D) Top: plot of cis-sister contact frequency P(s) as a function of genomic separation s for both sisters combined. Bottom: the derivative of the P(s) curve in the log-log space. (E) Aggregate cis-sister scsHi-C contact maps (±200 kb, log2 contact frequency normalized for distance-dependent decay) centered on contact enrichment "dots" detected in WT maps. Left - WT, right - NIPBL-AID. (F) Aggregate trans-sister scsHi-C contact maps (±500 kb) centered on stripes of enriched trans-sister contacts, detected in WT. Left - WT, right - NIPBL-AID. (G-J) Plots of cis- and trans-sister contact frequency P(s) as a function of genomic separation s for four conditions with perturbed cohesins - NIPBL-AID (G), Sororin-AID (H), prometaphase cells (I), and Rad21-AID (J). The gray lines show the three corresponding curves for "baseline" samples, WT in (G-I), and NIPBL-AID in (J). The arrows and numbers show the change in trans-contact frequency at 100 kb, compared to the "baseline" sample.

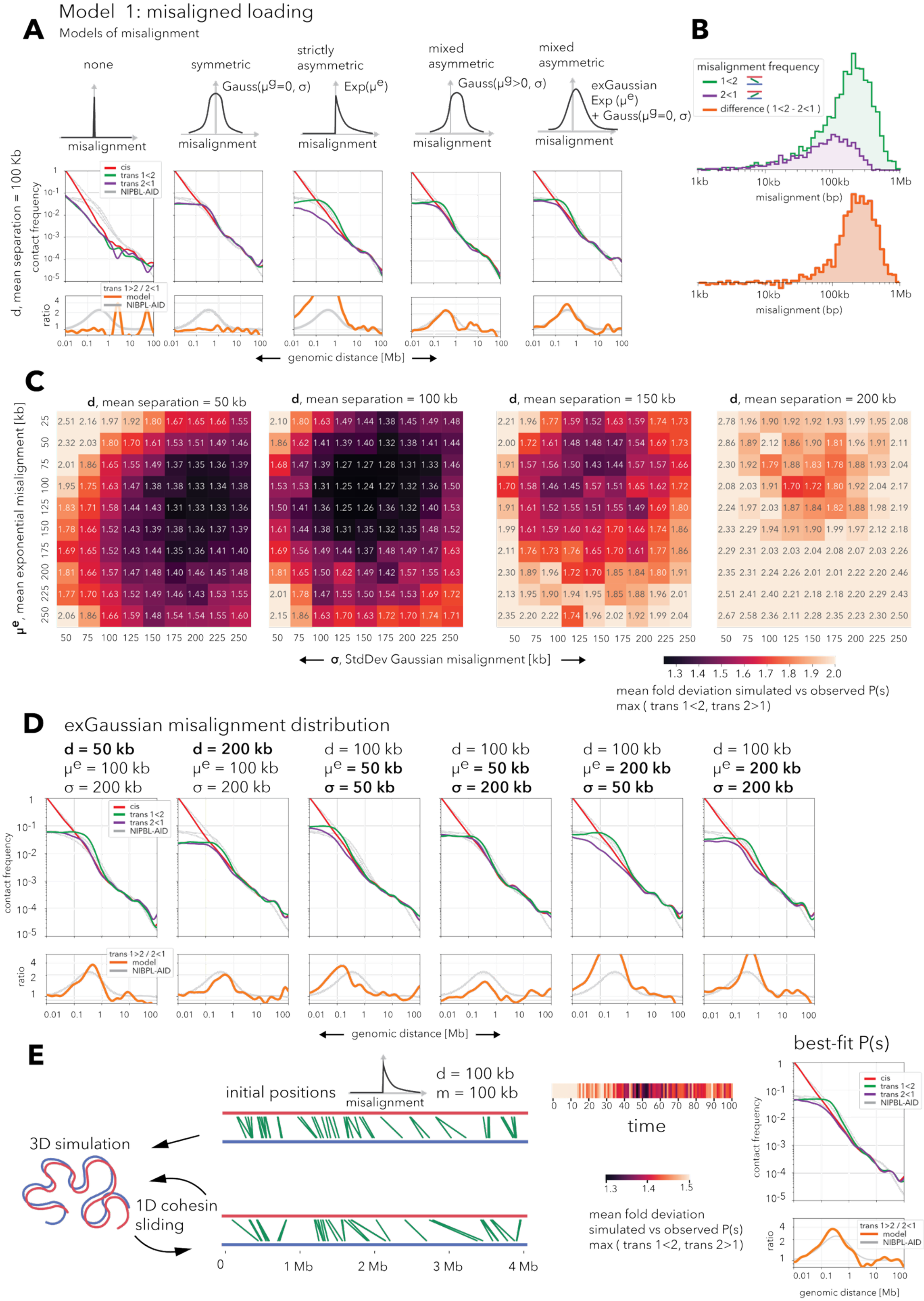

Model 1: misaligned loading
A
Models of misalignment
none
symmetric
strictly asymmetric
mixed asymmetric
mixed asymmetric
Gauss(μg=0, σ)
Exp(μe)
Gauss(μg>0, σ)
exGaussian Exp (μe) + Gauss(μg=0, σ)
misalignment
d, mean separation = 100 Kb
contact frequency
cis
trans 1<2
trans 2<1
NIPBL-AID
ratio
trans 1>2 / 2<1
model
NIPBL-AID
genomic distance [Mb]
B
misalignment frequency
1<2
2<1
difference ( 1<2 - 2<1 )
1kb
10kb
100kb
1Mb
misalignment (bp)
C
μe, mean exponential misalignment [kb]
d, mean separation = 50 kb
2.51 2.16 1.97 1.92 1.80 1.67 1.65 1.66 1.55
2.32 2.03 1.80 1.70 1.61 1.53 1.51 1.49 1.46
2.01 1.86 1.65 1.55 1.49 1.37 1.35 1.36 1.39
1.95 1.75 1.63 1.47 1.38 1.38 1.33 1.34 1.38
1.83 1.71 1.58 1.44 1.43 1.31 1.33 1.33 1.36
1.78 1.66 1.52 1.43 1.39 1.38 1.36 1.37 1.39
1.69 1.65 1.52 1.43 1.44 1.35 1.36 1.41 1.40
1.81 1.66 1.57 1.47 1.48 1.40 1.46 1.45 1.48
1.77 1.70 1.63 1.52 1.49 1.46 1.43 1.53 1.55
2.06 1.86 1.66 1.59 1.48 1.54 1.54 1.55 1.60
d, mean separation = 100 kb
2.10 1.80 1.63 1.49 1.44 1.38 1.45 1.49 1.48
1.86 1.62 1.41 1.39 1.40 1.32 1.38 1.41 1.44
1.68 1.47 1.39 1.27 1.27 1.28 1.31 1.33 1.46
1.53 1.53 1.31 1.25 1.24 1.27 1.32 1.36 1.50
1.50 1.41 1.36 1.25 1.26 1.36 1.32 1.40 1.53
1.61 1.44 1.38 1.31 1.35 1.32 1.35 1.48 1.52
1.69 1.56 1.49 1.44 1.37 1.48 1.49 1.47 1.63
1.81 1.65 1.50 1.62 1.49 1.42 1.57 1.55 1.63
2.01 1.78 1.47 1.55 1.50 1.54 1.53 1.69 1.72
2.15 1.86 1.63 1.70 1.63 1.72 1.70 1.74 1.71
d, mean separation = 150 kb
2.21 1.96 1.77 1.59 1.52 1.63 1.59 1.74 1.73
2.00 1.72 1.61 1.48 1.48 1.47 1.54 1.69 1.73
1.91 1.57 1.56 1.50 1.43 1.44 1.57 1.57 1.66
1.70 1.58 1.46 1.45 1.57 1.51 1.65 1.62 1.72
1.91 1.61 1.52 1.55 1.51 1.55 1.49 1.69 1.80
1.99 1.61 1.59 1.60 1.62 1.70 1.66 1.74 1.86
2.11 1.76 1.73 1.76 1.65 1.70 1.61 1.77 1.86
2.30 1.89 1.96 1.72 1.70 1.85 1.84 1.80 1.91
2.13 1.95 1.90 1.94 1.95 1.85 1.88 1.96 2.01
2.35 2.20 2.22 1.74 1.96 2.02 1.92 1.99 2.04
d, mean separation = 200 kb
2.78 1.96 1.90 1.92 1.92 1.93 1.99 1.95 2.08
2.86 1.89 2.12 1.86 1.90 1.81 1.96 1.91 2.11
2.30 1.92 1.79 1.88 1.83 1.78 1.88 1.93 2.04
2.08 2.03 1.91 1.70 1.72 1.80 1.92 2.04 2.17
2.24 1.97 2.03 1.87 1.84 1.82 1.88 1.98 2.19
2.33 2.29 1.94 1.91 1.90 1.99 1.97 2.12 2.18
2.29 2.31 2.03 2.03 2.04 2.08 2.07 2.03 2.26
2.35 2.31 2.24 2.26 2.01 2.01 2.22 2.20 2.46
2.51 2.41 2.40 2.21 2.18 2.18 2.20 2.29 2.43
2.67 2.58 2.36 2.11 2.28 2.35 2.23 2.30 2.50
σ, StdDev Gaussian misalignment [kb]
mean fold deviation simulated vs observed P(s)
max ( trans 1<2, trans 2>1)
D
exGaussian misalignment distribution
d = 50 kb μe = 100 kb σ = 200 kb
d = 200 kb μe = 100 kb σ = 200 kb
d = 100 kb μe = 50 kb σ = 50 kb
d = 100 kb μe = 50 kb σ = 200 kb
d = 100 kb μe = 200 kb σ = 50 kb
d = 100 kb μe = 200 kb σ = 200 kb
contact frequency
ratio
genomic distance [Mb]
E
initial positions
misalignment
d = 100 kb
m = 100 kb
3D simulation
1D cohesin sliding
0
1 Mb
2 Mb
3 Mb
4 Mb
time
mean fold deviation simulated vs observed P(s) max ( trans 1<2, trans 2>1)
best-fit P(s)
contact frequency
genomic distance [Mb]

**Figure S4. The model of misaligned cohesins quantitatively reproduces the trans-sister scsHi-C data.** (A) The effect of different misalignment distributions on trans P(s). Top: shapes of the tested misalignment distributions. Bottom: corresponding in silico P(s) curves and the log2 ratio of trans P(s) curves in the two shift directions (trans 1<2/trans 2<1) for one example parameter set of the corresponding distribution. (B) Log-scale histograms of cohesin misalignments in the two shift directions (top) and their difference (bottom) for the best-fitting parameter set (same data as in Fig. 4C). (C) Heatmaps showing the mean fold deviation between simulated and observed trans P(s) (maximum of trans 1<2 and trans 2<1 deviations) for the tested parameters of cohesin distribution. Three parameters were systematically varied: the average cohesin separation, the mean of the exponential component, and the standard deviation (sigma) of the Gaussian component of the ex-Gaussian distribution of cohesin misalignment. (D) Effects of parameter variations on simulated trans P(s). Variation in the three parameters (average cohesin separation d, mean $\mu^e$ of the exponential component, and standard deviation $\sigma$ of the Gaussian component of the exGaussian misalignment distribution) affects the shape of trans P(s). Increasing average separation d (i.e., decreasing cohesin frequency) shifts the trans curve downward (compare panels 1 and 2). Increasing the standard deviation $\sigma$ of non-directional (i.e. Gaussian) misalignment closes the gap between the two trans curves (compare panels 3 vs 4 and 5 vs 6). Increasing the mean $\mu^e$ of the directional (exponential) misalignment widens the gap between the two trans P(s) curves and extends it towards longer separations. (E) The model of freely sliding cohesins, initially misaligned strictly asymmetrically. Left: the illustration of the approach. Cohesin spacings and misalignments are both initially sampled from an exponential distribution (d = 100 kb, $\mu^e$ = 100 kb). The simulation proceeds through alternating steps in 3D (modeling diffusion of chromatin) and 1D (modeling sliding of cohesins along chromatids). Middle: the heatmap of simulated-vs-observed P(s) discrepancy for sequential steps of a simulation. Right: the simulated P(s) for the best-fit step of the simulation.

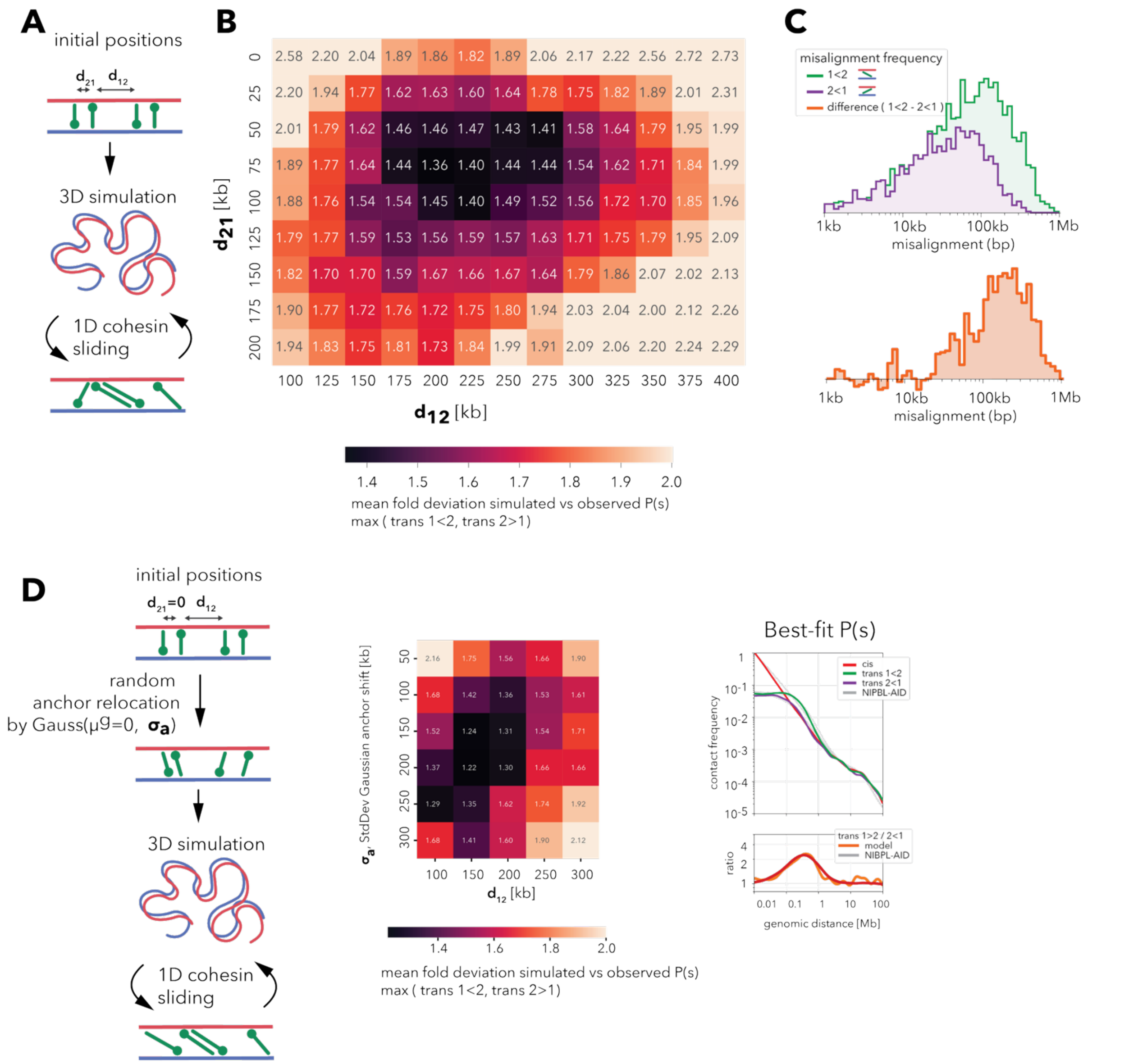


**Figure S5. The model of paired semi-anchored cohesion can quantitatively fit the trans-sister scsHi-C data.** (A) The schematics of the simulation of paired semi-anchored cohesion. Sequential cohesins are semi-anchored to different sisters in an alternating order. The spacings between resulting "2-1" and "1-2" pairs are sampled from two different exponential distributions with the means of $d_{21}$ and $d_{12}$. The simulation proceeds through alternating steps in 3D (modeling diffusion of chromatin) and 1D (modeling sliding of semi-anchored cohesins along chromatids). (B) The heatmap of the mean fold deviation between simulated and observed trans P(s) (maximum of trans 1<2 and trans 2<1 deviations) as a function of $d_{21}$ and $d_{12}$ cohesin spacings. (C) Log-scale histograms of cohesin misalignments in the two shift directions (top) and their difference (bottom) for the best-fitting parameter set (same data as in Fig. 5F). (D) A modification of the "paired semi-anchored" model, where cohesins are placed in tight "2-1" pairs ($d_{21}$=0), but their anchors are displaced randomly by external processes. Left: the flow of the simulation is the same as in (A), except that, after initialization, the positions of the anchors are perturbed randomly by a Gaussian-distributed amount with the standard deviation σ. Middle: the heatmap of simulated-vs-observed P(s) discrepancy for the two model parameters, $d_{12}$ and σ. Right: the simulated P(s) for the best-fit set of parameters.

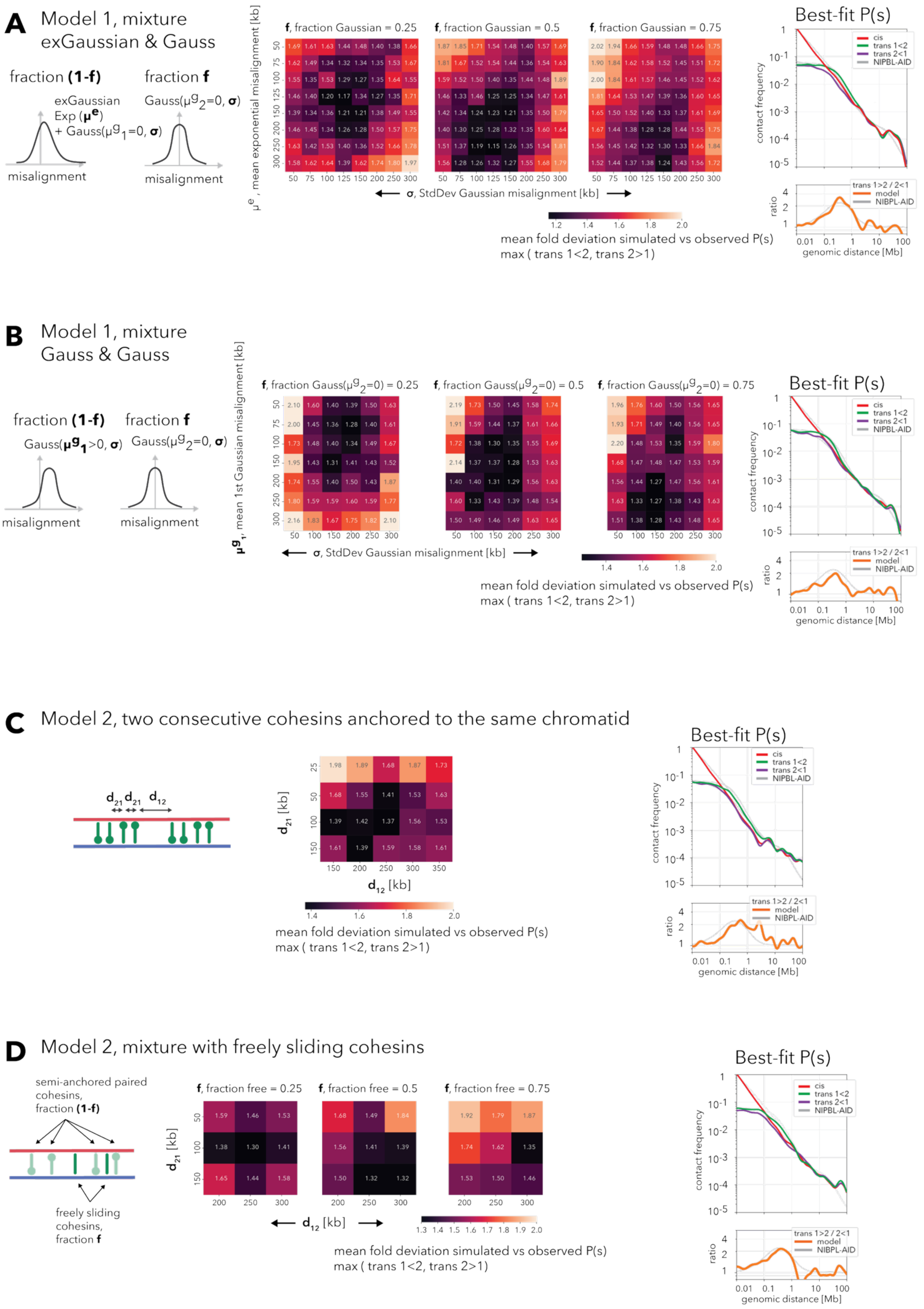

A
Model 1, mixture exGaussian & Gauss
fraction (1-f)
fraction f
exGaussian Exp (μe) + Gauss(μg1=0, σ)
Gauss(μg2=0, σ)
misalignment
misalignment
μe, mean exponential misalignment [kb]
f, fraction Gaussian = 0.25
f, fraction Gaussian = 0.5
f, fraction Gaussian = 0.75
σ, StdDev Gaussian misalignment [kb]
mean fold deviation simulated vs observed P(s)
max ( trans 1<2, trans 2>1)
Best-fit P(s)
contact frequency
cis
trans 1<2
trans 2<1
NIPBL-AID
ratio
trans 1>2 / 2<1
model
NIPBL-AID
genomic distance [Mb]
B
Model 1, mixture Gauss & Gauss
fraction (1-f)
fraction f
Gauss(μg1>0, σ)
Gauss(μg2=0, σ)
misalignment
misalignment
μg1, mean 1st Gaussian misalignment [kb]
f, fraction Gauss(μg2=0) = 0.25
f, fraction Gauss(μg2=0) = 0.5
f, fraction Gauss(μg2=0) = 0.75
σ, StdDev Gaussian misalignment [kb]
mean fold deviation simulated vs observed P(s)
max ( trans 1<2, trans 2>1)
Best-fit P(s)
contact frequency
ratio
genomic distance [Mb]
C
Model 2, two consecutive cohesins anchored to the same chromatid
d21 d21 d12
d21 [kb]
d12 [kb]
mean fold deviation simulated vs observed P(s)
max ( trans 1<2, trans 2>1)
Best-fit P(s)
contact frequency
ratio
genomic distance [Mb]
D
Model 2, mixture with freely sliding cohesins
semi-anchored paired cohesins, fraction (1-f)
freely sliding cohesins, fraction f
f, fraction free = 0.25
f, fraction free = 0.5
f, fraction free = 0.75
d21 [kb]
d12 [kb]
mean fold deviation simulated vs observed P(s)
max ( trans 1<2, trans 2>1)
Best-fit P(s)
contact frequency
ratio
genomic distance [Mb]

**Figure S6. Models allow the existence of multiple cohesin populations with different misalignment statistics.** (A-B) Modifications of model 1, where cohesins come from two subpopulations with different misalignment statistics: (A) asymmetric exGaussian and symmetric Gaussian (mean=0) and (B) asymmetric Gaussian (mean > 0) and symmetric Gaussian (mean=0). Left: cartoon illustration of the misalignment distributions of the two subpopulations of cohesins. Middle: the heatmap of the mean fold deviation between simulated and observed trans P(s) (maximum of trans 1<2 and trans 2<1 deviations) as a function of the fractions and parameters of the two mixed distributions. Right: the simulated P(s) for the best-fit mixture model. (C-D) Modifications of model 2: (C) two consecutive cohesins are anchored to the same sister chromatid, and (D) cohesins have two subpopulations: "paired semi-anchored" and freely sliding ones. Left: cartoon illustration of the model. Middle: the heatmap of the mean fold deviation between simulated and observed trans P(s) (maximum of trans 1<2 and trans 2<1 deviations) as a function of the model parameters. Right: the simulated P(s) for the best-fit mixture model.

9. G. Wutz, R. Ladurner, B. G. St Hilaire, R. R. Stocsits, K. Nagasaka, B. Pignard, A. Sanborn, W. Tang, C. Várnai, M. P. Ivanov, S. Schoenfelder, P. van der Lelij, X. Huang, G. Dürnberger, E. Roitinger, K. Mechtler, I. F. Davidson, P. Fraser, E. Lieberman-Aiden, J.-M. Peters, ESCO1 and CTCF enable formation of long chromatin loops by protecting cohesinSTAG1 from WAPL. *Elife* **9**

(2020).
